# A robust workflow to benchmark deconvolution of multi-omic data

**DOI:** 10.1101/2024.11.08.622633

**Authors:** Elise Amblard, Vadim Bertrand, Luis Martin Pena, Slim Karkar, Florent Chuffart, Mira Ayadi, Aurélia Baurès, Lucile Armenoult, Yasmina Kermezli, Jérôme Cros, Yuna Blum, Magali Richard

## Abstract

Tumour heterogeneity significantly affects cancer progression and therapeutic response, yet quantifying it from bulk molecular data remains challenging. Deconvolution algorithms, which estimate cell-type proportions in bulk samples, offer a potential solution. However, there is no consensus on the optimal algorithm for transcriptomic or methylomic data. Here, we present an unbiased evaluation framework for the first comprehensive comparison of deconvolution algorithms across both omic types, including reference-based and -free approaches. Our evaluation covers raw performance, stability, and computational efficiency under varying conditions, such as missing or additional cell types and diverse sample compositions. We apply this framework across multiple benchmark datasets, including a novel multi-omics dataset generated specifically for this study. To ensure transparency and re-usability, we have designed a reproducible workflow using containerization and publicly available code. Our results highlight the strengths and limitations of various algorithms, and provides practical guidance for selecting the best method based on data type and analysis context. This benchmark sets a new standard for evaluating deconvolution methods and analysing tumour heterogeneity.

## Introduction

Bulk transcriptome and methylome are routinely measured in the clinic to diagnose and classify cancer patients. However, these data are usually analysed in a way that do not account for intra-tumour heterogeneity, i.e. the fact that a tumour sample is composed of different cell types [1]. However, intra-tumour heterogeneity is a critical parameter as it will have an impact on the tumour evolution and its response to treatment [2, 3]. It is possible to extract this information from bulk data with deconvolution tools that aim at predicting the proportions of the different cell types present in the sample [1, 4]. Deconvolution can also be used to quantify heterogeneity in non-cancer data, but deconvolving cancer tissues is the most classical use case in the literature. Still, there is no strong consensus on the best method to use [5, 6]. Secondly, as there is no multi-omic benchmark to our knowledge, it is not known what is the easiest omic to analyse with respect to the deconvolution task.

There are two classes of deconvolution algorithms. The supervised (or reference-based) methods use a matrix of reference profiles to estimate cell-type proportions, while the unsupervised (or reference-free) methods estimate simultaneously the proportions along with reference molecular profiles of each cell type. In the supervised class, the quality of the references is key for the deconvolution performance [7, 8]. Critical points include the fact that the cells in the references should preferably come from the same tissue context as those to be deconvoluted (*in vivo* or *in vitro*), and that it should contain expected cell types [6, 7, 9]. More precisely, most supervised algorithms cannot handle missing cell types. One solution implemented by few methods, called semi-supervised methods, is to allow the prediction of an unknown component [10]. On the other hand, unsupervised methods do not rely on references, avoiding associated challenges, while major disadvantages have been the higher complexity of the problem and the difficulty to identify the cell types retrieved during deconvolution. As a matter of fact, it has been shown that the components obtained after deconvolution are likely a linear combination of cell types present in the samples [2, 11].

Single-cell based methods would allow to overcome deconvolution-related issues by allowing a straightforward quantification of intra-tumour heterogeneity. Nonetheless, integrating such technologies into the standard of care for hospital patients is not yet feasible due to their time-consuming nature and high costs. It was also observed that single-cell technologies do not capture all cell types with the same probability, blurring the estimation of the cellular composition [12]. Conversely, bulk technologies have been used for a long time at the hospital, and focusing on the analysis of these data will also allow to leverage the massive amount of data already generated. Nonetheless, it is possible to take advantage of both single-cell and bulk technologies. The strategy is to sequence the single cells of few samples to generate reference profiles from the same context and use them for supervised deconvolution of the other samples. We did not include single-cell-based deconvolution methods, while they are more recent than bulk-based ones, for two main reasons. First, the properties of a bulk transcriptome differ significantly from those of a single-cell transcriptome, owing to variations in tissue processing and RNA enrichment protocols [13]. As a result, using single-cell references can be misleading compared to bulk references. Indeed, a previous benchmark showed that second generation single-cell-based methods did not outperform state-of-the-art bulk-based tools [14]. Second, there is no single-cell-based DNAm deconvolution methods. However, second generation single-cell-based methods are powerful and will probably be one of the lead to improve deconvolution methods [14, 15]. Another lead is to leverage bulk multi-omic approaches, which is the motivation of this benchmark.

As of today, there is a humongous amount of deconvolution algorithms available. Many benchmarks were published to help bioinformaticians choose the best tool as a function of their data (Table 1). Still, current benchmarks suffer from several pitfalls: most of them include less than 10 methods, they do not all confirm rankings on real-world datasets (termed silver-standard datasets) and none except one provides a single comprehensive ranking. Only few looked at supervised and unsupervised methods simultaneously, and all of them studied a single omic: either transcriptomic or methylation alone. Lastly, there is a high discrepancy between these benchmarks, one possible reason being the inconsistency of the metrics used to measure performance [6].

**Table 1.** Characteristics of selected benchmark studies. AE: Absolute Error. RMSE: Root Mean Square Error. MAE: Mean Absolute Error. MAPE: Mean Absolute Percentage Error. sMAPE: symmetric MAPE. mAD: mean Absolute Deviation.

| Benchmark ID | Modality | Methods (#) | Class | Metrics | Gold-standards | Silver-standards |
| --- | --- | --- | --- | --- | --- | --- |
| [7] | RNA | **20** | **Both** | RMSE, Pearson correlation | ✓ | ✓ |
| [8] | RNA | 12 | Supervised | AE, Pearson correlation | ✓ | ✓ |
| [16] | RNA | 3 | Supervised | RMSE, MAE, MAPE, sMAPE, Pearson correlation | ✓ | ✓ |
| [17] | RNA | 11 | **Both** | mAD, Pearson correlation | ✓ | ✗ |
| [18] | RNA | 7 | Supervised | Pearson correlation | ✓ | ✓ |
| Our framework | **RNA** | 16 | **Both** | RMSE, MAE, Pearson correlation,<br>**Overall comprehensive score** | ✓ | ✓ |
| [19] | DNAm | 3 | Unsupervised | RMSE, MAE, Pearson correlation | ✓ | ✓ |
| [20] | DNAm | 4 | Supervised | RMSE, Pearson correlation | ✓ | ✓ |
| [21] | DNAm | 8 | **Both** | RMSE, sMAPE, Spearman correlation | ✓ | ✗ |
| Our framework | **DNAm** | 11 | **Both** | RMSE, MAE, Pearson correlation,<br>**Overall comprehensive score** | ✓ | ✓ |

In our benchmark, we addressed those pitfalls. We built a robust workflow to systematically rank and evaluate deconvolution methods. We included 20 algorithms, supervised or unsupervised, designed for transcriptome or methylome data. We tested all methods on 3 types of data: *in silico* simulations, *in vitro* mixes and *in vivo* data, including a new multi-omic dataset. We studied various aspects of a method’s performance: (i) raw performance which measures errors in the prediction of the proportions, (ii) stability, and (iii) execution time. Based on those 3 categories, we provide an overall score per method along with intermediate scores, and p-values. We evaluated the effect of missing or extra cell types in supervised deconvolution, and of data dispersion and size in unsupervised deconvolution. We also analysed the methods’ performance in terms of rare cell types detection. We related our ranking with findings from previous benchmarks. We provide (i) a GitHub repository with all the codes needed to reproduce the analysis performed in the paper, along with (ii) an Apptainer container to run all methods included, and (iii) guidelines to choose a deconvolution method among those included here, as a function of the data to analyse. The repository and the container have been designed with flexibility in mind, allowing to include new datasets and methods.

## Results

Our benchmark is a combination of two pipelines (Figure 1A). The first pipeline performs the deconvolution task, the second the ranking task. The ranking pipeline has been designed to compare extensively deconvolution algorithms.

**Fig. 1.**
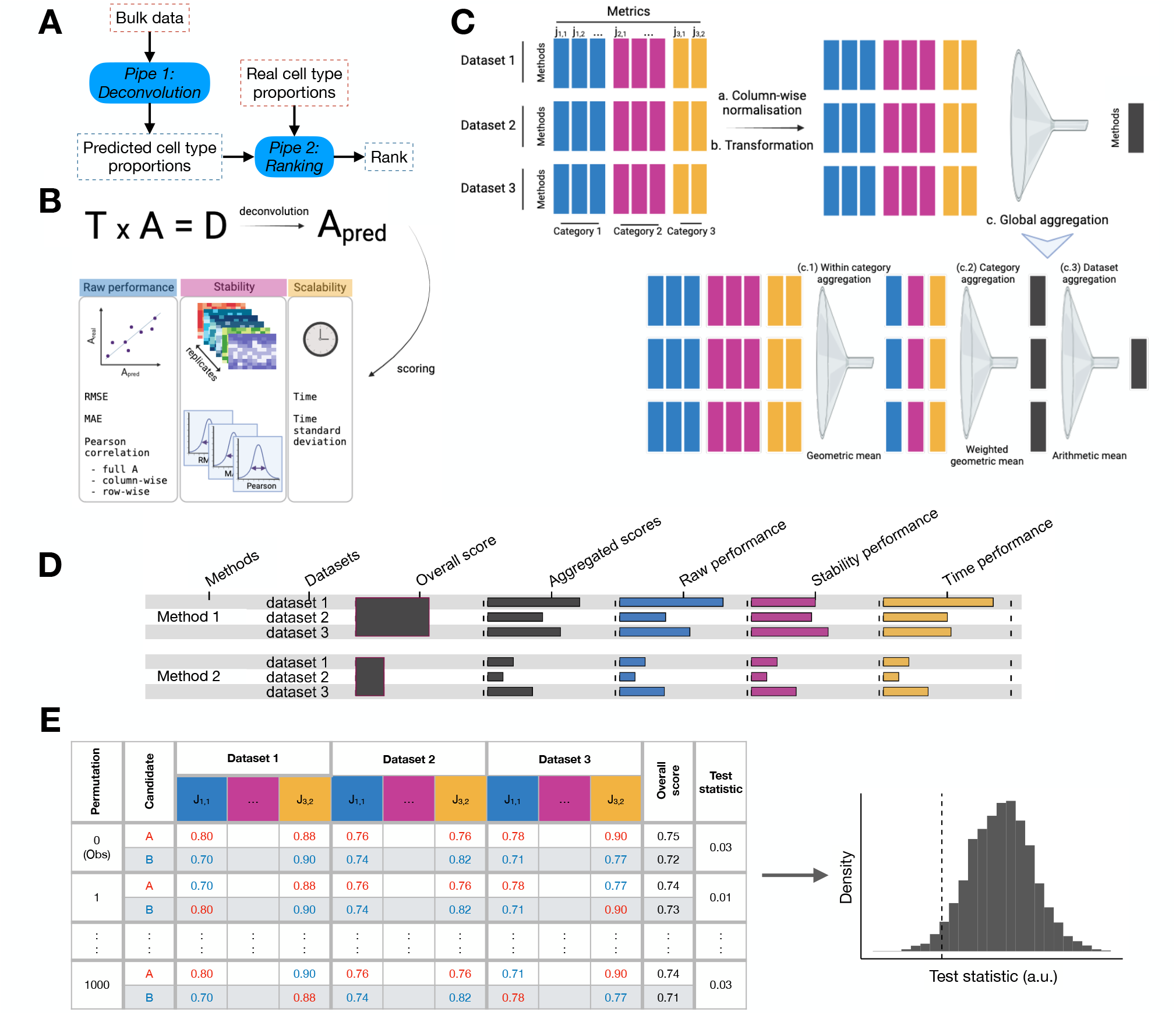
A comprehensive benchmark pipeline evaluates several aspects of the deconvolution performance. **(A)** The pipeline is done in 2 steps, a deconvolution step and a ranking step. **(B)** The deconvolution step outputs an estimation of the proportions which is used to evaluate a method’s raw performance and stability, along with its running time. **(C)** Metrics aggregation procedure, recapitulating a series of metrics from different natures across several datasets and yielding a single overall score per method. The different natures of the metrics imposes to first normalize and transform the scores such that they all lie between 0 and 1, 1 being the best possible value. **(D)** Example of an output from the 3-steps global aggregation procedure with the display of intermediate scores. **(E)** Permutation test to compute p-values.

### The deconvolution pipeline is exhaustive, reproducible and flexible

We simulated 6 datasets (3 multi-omic, 2 RNA, 1 DNAm datasets), termed gold-standard, based on the bulk molecular profiles of pure cell types (Table 2). Using simulations allowed us to easily evaluate the performance of deconvolution methods as the ground truth is readily available from the simulation process. On the contrary, real datasets’ ground truth is usually not known, or only accessible via proxies such as FACS or imaging. Another advantage of simulated datasets is the possibility to test the impact of data characteristics on the quality of the deconvolution. Here, we tested the effect of the number of samples (*n* = 30, 120) and of the diversity in sample composition with three levels of dispersion. Despite those benefits, the drawback of using simulations is that the results of the benchmark might not comply with real-life scenarii. Hence, we also analysed real datasets to confront our rankings (Table 2). We added gold-standard *in vitro* mixes: 1 multi-omic, 1 DNAm and 1 RNA datasets; as well as 2 RNA and 2 DNAm silver-standard *in vivo* datasets. *In vitro* mixes are real-life datasets, with the advantage of having the ground truth, but there are only few such datasets available.

**Table 2.**
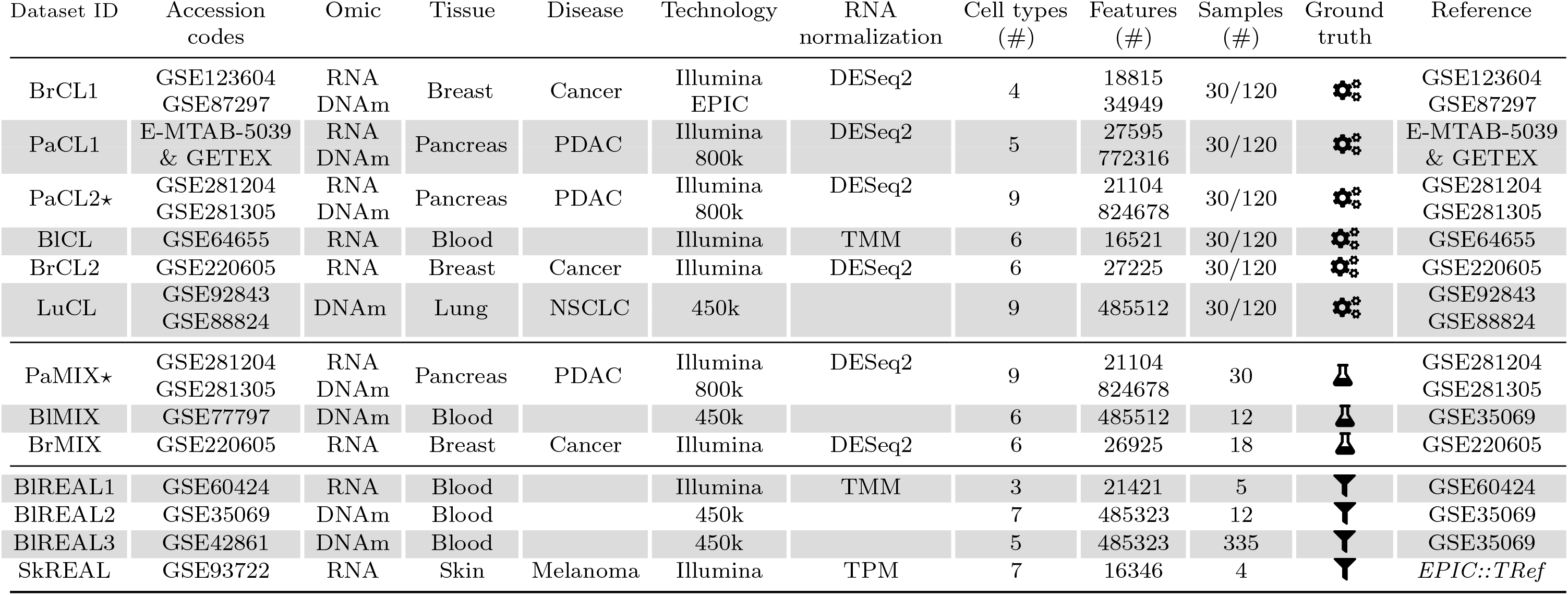
Table of gold- and silver-standard datasets used in the benchmark. The penultimate column refers to the type of the ground truth matrix: for gold-standard datasets, the ground truth is exact and represented by the symbols 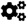 for simulations and 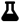 for the *in vitro* mixes, while it is a proxy, usually measured by FACS and represented by the symbol 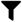, for silver-standard datasets. The last column refers to the reference profiles that were used for supervised deconvolution. The *in silico* datasets are under the acronym CL (Cell Line), the *in vitro* datasets under the acronym MIX and the *in vivo* under the acronym REAL. New datasets generated for this study are flagged with the symbol ⋆. For the PaCL1 dataset, more details on how to download it are on the website https://deconbench.github.io/Benchmarkdatasets.html.

We analysed thirteen gold and silver-standard datasets (Table 2) with twenty deconvolution methods (see Methods and Supplementary Tables 1, 2) in the deconvolution task. All methods have been run on a high performance computing cluster, using Snakemake [22] and an Apptainer container [23] (see Methods) to enforce reproducibility and re-usability. Thanks to these tools, the deconvolution pipeline is flexible in order to allow for additional methods and/or datasets. The input of this first pipeline is a dataset and a method and the output is the estimation of cell type proportions along with the running time (Figure 1A,B).

We tested several deconvolution methods designed for either of the two omics we included, namely the methylome and transcriptome, comprising different algorithmic designs: Bayesian-, or LS-based (Least Squares) for supervised methods, ICA- or NMF-based for unsupervised ones. After a thorough review of the literature (Supplementary Tables 1, 2), we included 12 supervised methods in our deconvolution pipeline (n=7 for transcriptomic, n=1 for methylation data, n=4 for both omics), and 8 unsupervised methods (n=2 for transcriptomic, n=3 for methylation data, n=3 for both omics) (see Methods, paragraphs “Supervised deconvolution algorithms” and “Unsupervised deconvolution algorithms” for an exhaustive list of the algorithms we tested).

### The ranking process enables unambiguous comparisons of algorithms

The ranking pipeline takes as input the estimation of the proportions and the running time from the deconvolution pipeline. The output of this second pipeline is an overall benchmark score associated to p-values for the pairwise comparisons of methods, along with the intermediate scores (Figure 1A,E). For each prediction, we computed five performance metrics along with the running time, named primary metrics, and six additional secondary metrics when replicates were available (Figure 1B and Methods): error metrics (RMSE, MAE and Pearson correlations) to measure the raw performance, standard deviation of those error metrics to measure the stability, and the running time and its stability. We selected metrics that are routinely used in other benchmarks (Table 1). Those metrics are aggregated by the pipeline into a single overall score (Figure 1C and Methods). We also display intermediate scores in order to better understand why a method has a good or bad performance (Figure 1D). Finally, we compute p-values along with the overall score (Figure 1E and Methods).

We designed several ranking processes and tested them according to empirical criteria defined in [24] (Supplementary Figure 1 and Methods). We chose the process *S*_*consensus*_, as it was the most generalizable one, based on the generalisation criterion. This process averages the output of three other processes *S*_*raw*_, *S*_*rank*_ and *S*_*topsis*_. *S*_*raw*_ merges the different metrics after normalization, *S*_*rank*_ merges the ranks computed from each metric, and *S*_*topsis*_ merges the TOPSIS scores [25] (see Methods for more details). This criterion quantifies the stability of the ranking with respect to the exclusion of few metrics, which is a desired property. Besides, we observed that the top methods elected by each process remained the same, which serves as a proof of the robustness of our ranking processes and their results (Supplementary Figure 2). The codes to run the ranking process are available and can be modified to add new metrics or modify the aggregation process (See Methods).

### RLR is the best supervised method irrespective of the type of omic

We first analysed simulated data and applied deconvolution on 3 versions of each expression/methylation matrix: (i) without feature selection, with feature selection performed (ii) by selecting the 1,000 most variable features or (iii) by TOAST and selecting 1,000 features [26]. TOAST works by iteratively doing ICA followed by a selection of the features differentially expressed across components. For each deconvolution method, we selected the version of the expression matrix yielding the best overall benchmark score (Supplementary Figure 3). Overall, the strategy with no feature selection is the best ranked in 63% (17/27) of the cases, and it is especially true for RNA data (11/16). The second best feature selection task is TOAST (7/10). We observed that increased performance after feature selection is partly due to a substantially decreased running time, while stability is reduced (Supplementary Figure 4).

Figure 2 presents the result of the benchmark for supervised methods on simulated data, displaying the p-values along with overall and aggregated scores. RLR is one of the 4 methods that can be used on both omic types and it has been consistently the best method in a significant fashion (Figure 2A,B). RLR is also the method yielding the less variation in performance from one dataset to another (Supplementary Figure 5A,C). Those good results can be explained by the fact that the method is robust to outliers, while other least-squares-based methods have also good scores. On the contrary, the performance dependency with respect to the dataset increases with decreasing overall scores, especially for the methylome. For all datasets, RLR displays the best aggregated scores (Supplementary Figure 6A,C) as well as the best scores for the raw performance (as shown also in [27] for DNAm and in [7] for RNA) and the stability categories (Figure 2C,D). Additionally, our results are coherent with the ranking obtained in [27] where the authors used Spearman correlation.

**Fig. 2.**
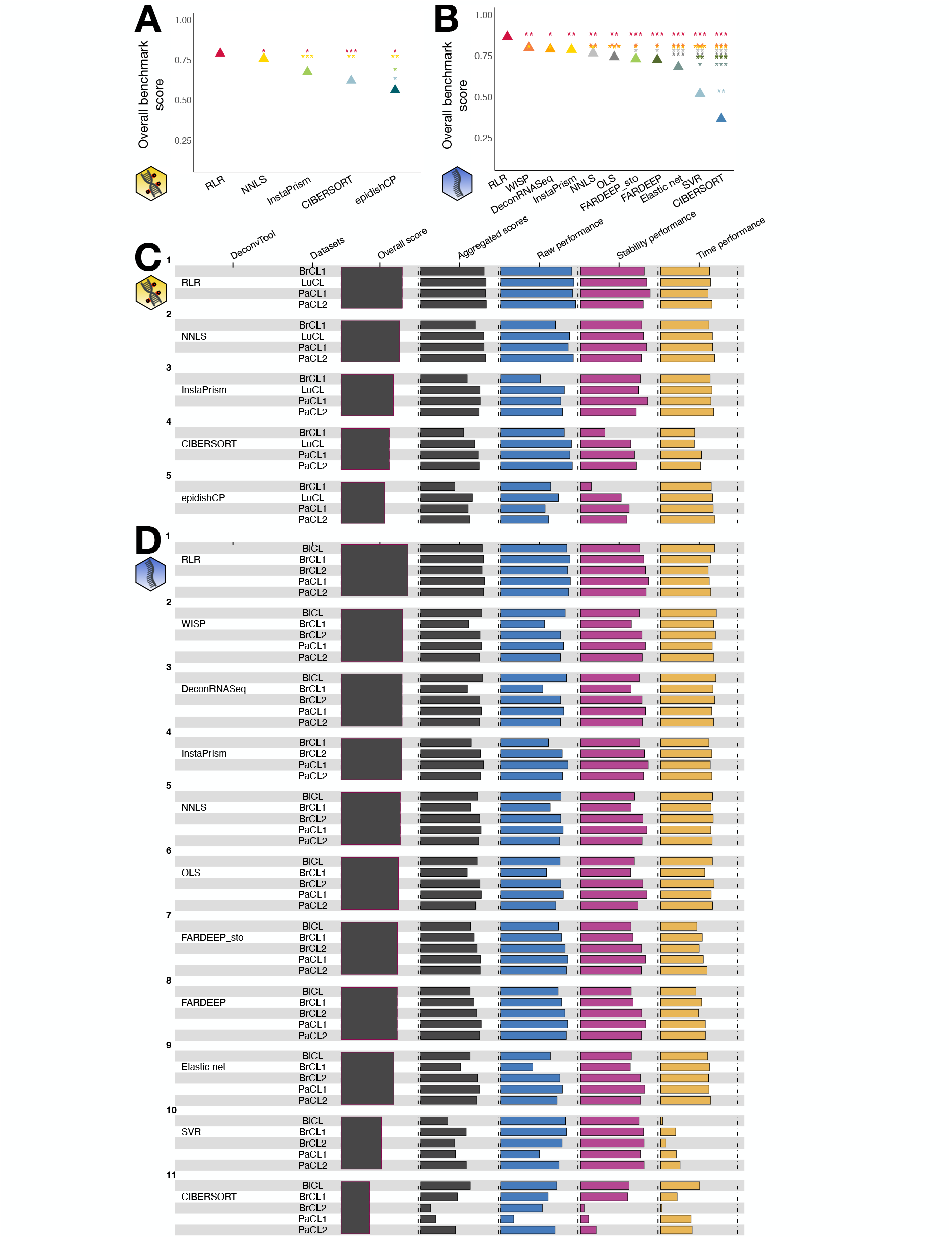
Benchmark scores of supervised deconvolution methods. Methods have been applied to the methylation **(A**,**C)** or the transcriptome **(B**,**D). (A**,**B)** Statistical significance of the differences in overall benchmark scores. Overall benchmark scores are shown with triangle symbols, and one-sided p-values between two methods are shown with stars, the colour and y-value relating to the supposedly better method, the x-value relating to the supposedly worse method. Stars follow the convention: (*) for p-values below 0.05, (**) for p-values below 0.01, (***) for p-values below 0.001.**(C**,**D)** Overall and intermediate scores. Each row represents a method run on the different simulations. The third column is the overall benchmark score, the forth the aggregated score per dataset, and the fifth to seventh columns the scores relating to each of the three categories, averaged per dataset. The yellow DNA icon stands for methylation data, the blue single-strand one for transcriptome data.

### The performance of unsupervised methods strongly depends on the dataset

While there is a consensus on the best method for both omics and across datasets in the supervised setting, it is not the case for unsupervised algorithms. Again, we retained the feature selection strategy which gave the best overall score for each deconvolution algorithm (Supplementary Figure 3). For methylation data, ICA has the highest overall score, and it does significantly better than all other methods (Figure 3A,C), but it displays contrasted results across datasets: it is the best algorithm for 50% of the datasets (2/4), and the second best for PaCL2 (Supplementary Figure 6B). On the contrary, the best methods are debCAM for PaCL1 and MeDeCom for PaCL2, and the method yielding the most stable results across datasets is EDec (Supplementary Figure 5B).

**Fig. 3.**
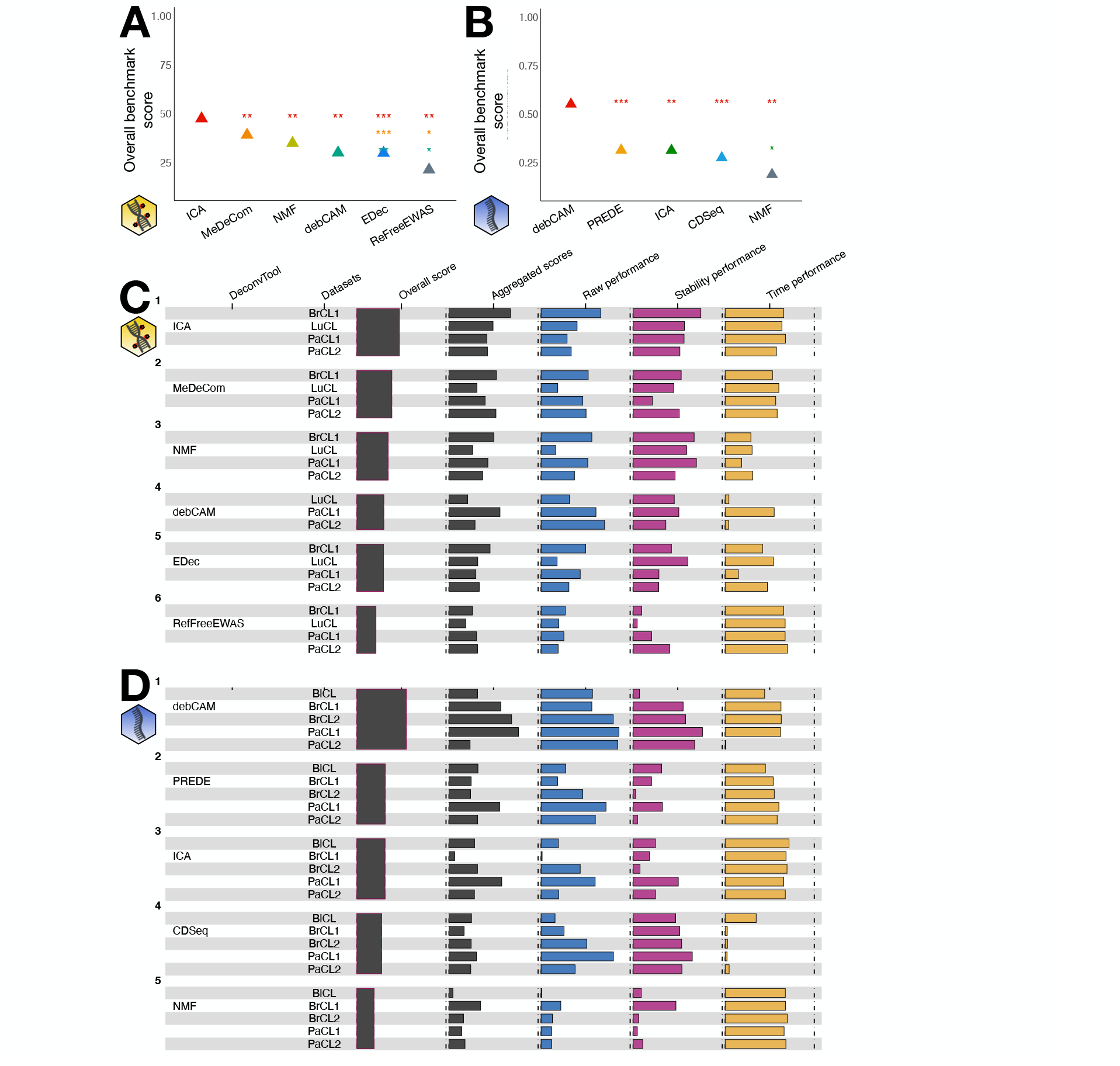
Benchmark scores of unsupervised deconvolution methods. Methods have been applied to the methylation **(A**,**C)** or the transcriptome **(B**,**D). (A**,**B)** Statistical significance of the differences in overall benchmark scores.**(C**,**D)** Overall and intermediate scores.

For transcriptomic data, the best method is debCAM (Figure 3B,D) as it performs particularly well in all except the PaCL2 dataset compared to the other methods (Supplementary Figure 6D). Again, the performance varies a lot from one dataset to another, except for ICA (Supplementary Figure 5D). We observed that the performance of unsupervised deconvolution depends on the dataset in a much stronger way than for supervised methods (Supplementary Figure 5). All those observations shed light on the difficulty of reaching a consensus on the best deconvolution tool. Additionally, supervised methods perform better than unsupervised ones as already shown in the literature [28, 29].

### RNA yields the best performance based on analysis of both types of omic

Since we have an interest in using multi-omic data, we wanted to determine the easiest omic to deconvolve (if any), based on the three simulated multi-omic datasets. We observed that whatever the class of the method, RNA was the omic with the highest overall benchmark score (Figure 4A). For the supervised class, RLR done on RNA data is the best option, and the second best choice is to use RLR on DNAm data. The next best methods are linked to RNA deconvolution. For the 4 methods that can be applied on both omics, three are optimal when used on RNA data: RLR, InstaPrism and NNLS. Regarding the unsupervised class, we also observed that the omic yielding the best performance is RNA analysed with debCAM, though in this class the next best methods are linked to DNAm analysis: ICA, MeDeCom and NMF. The three methods with which we can analyse both omics perform best when used on DNAm except for debCAM, which was designed for RNA analysis initially. Based on the datasets we analysed and the methods included, we would recommend to use RNA for cell-type quantification in case of a simple-omic supervised deconvolution. For the unsupervised deconvolution, the hegemony of the transcriptome is less marked, with several methylation methods having good results.

**Fig. 4.**
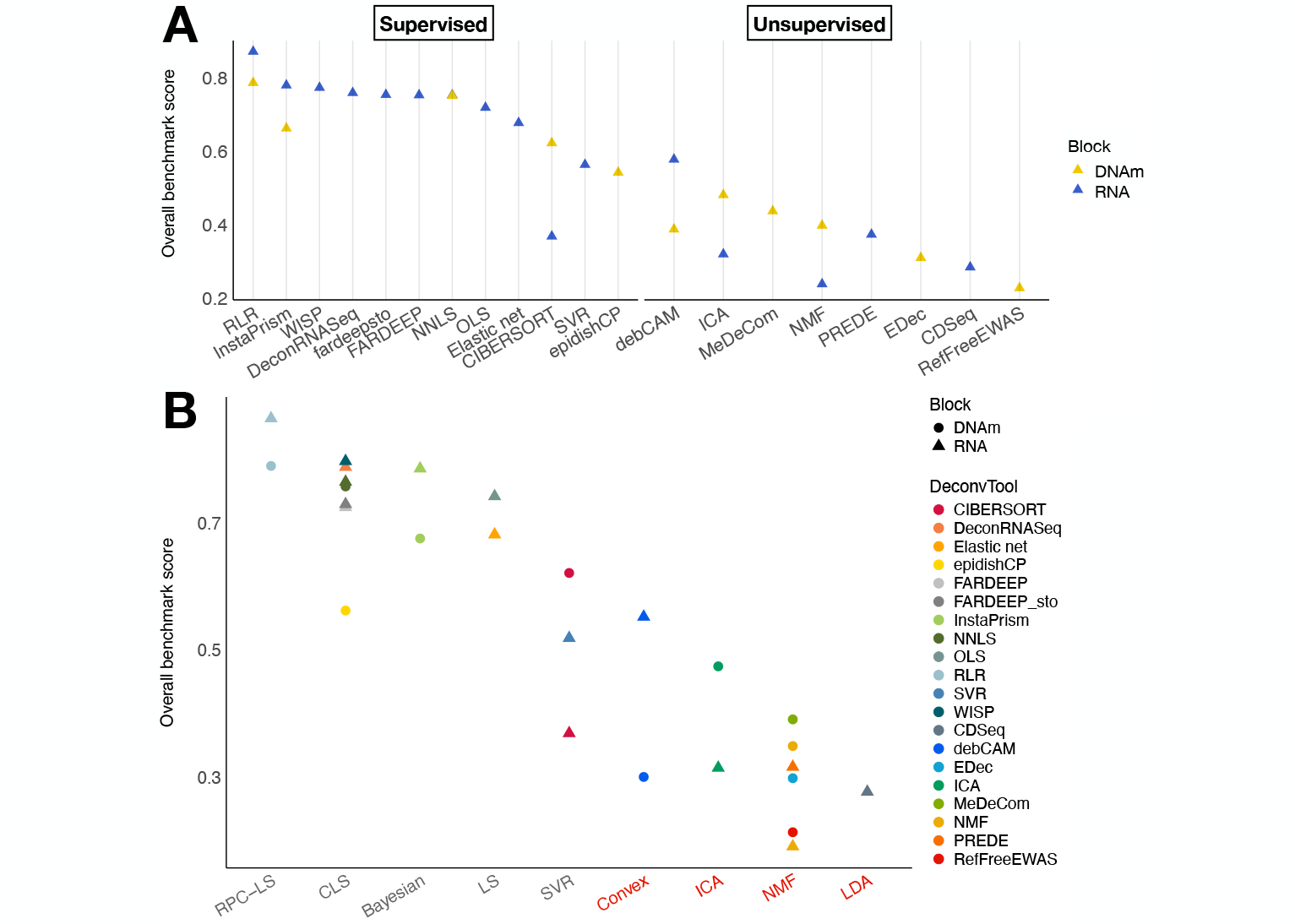
Comparison of the deconvolution performance for each omic. **(A)** Overall benchmark score of all methods included in the benchmark, ranked on the three multi-omic *in silico* datasets. **(B)** Overall benchmark scores of all methods sorted by design (*i*.*e*. type of algorithm), with unsupervised designs labelled in red. LS: Least Squares. CLS: Constrained LS. RPC: Robust Partial Correlation. SVR: Support Vector Regression. LDA: Latent Dirichlet Allocation.

We wanted to understand whether RNA deconvolution was more efficient because of a better algorithmic design or because of the intrinsic nature of the two omics we analysed. We tested what specific designs were the most competitive in our benchmark. We summarized in Supplementary Figure 7 the type of model, and whether the sum-to-one constraint on the proportion matrix was implemented in the form or an equality or inequality. Since there are too few methods that used an inequality constraint, we focused on analysing the impact of the type of model. The most popular approaches are NMF for the unsupervised class and LS for the supervised class (Supplementary Figure 7). However, the most efficient ones were the convex analysis and ICA for unsupervised methods and robust LS for supervised methods (Figure 4B). While it provides hints for future algorithms to improve the quality of the deconvolution, we also observed that for a given design applied to both omics, the best benchmark score is attributed to RNA in 50% of the cases (3/6). In particular, LS-based methods perform best on RNA data, but NMF-based methods perform best on DNAm data. To conclude on this, each approach seems to perform best on a specific omic, probably relating to the property of the data itself.

### Simulation parameters have an impact on the deconvolution performance

We tested two simulation parameters, namely the number of samples (*n* = 30, 120) and the dispersion factor (*α*_0_ = 3, 10, 30), related to the amount of heterogeneity in the proportions: the higher *α*_0_, the lower the dispersion (see Methods). We considered the baseline simulation as the one defined by *n* = 120, *α*_0_ = 10, and we evaluated the impact on the overall score of varying the number of samples or the dispersion, mimicking biological noise, in the unsupervised class of methods.

For the number of samples, we observed that unsupervised methods always perform worse whenever there are less samples than in the baseline (Figure 5A), especially for the methylation where score differences are significant for all methods except one. This trend is expected and has been shown before [19].

**Fig. 5.**
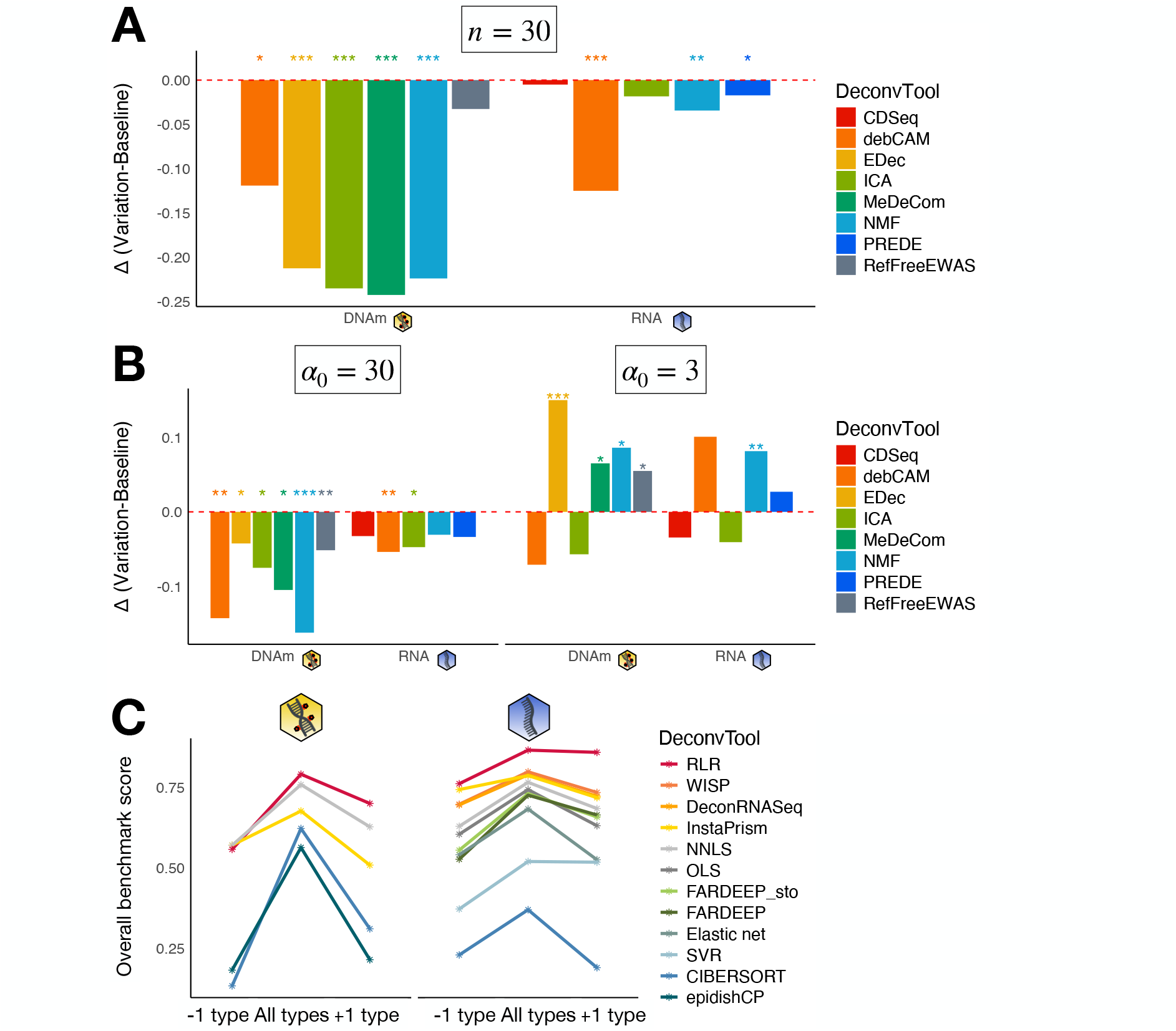
Impact of simulation parameters. **(A)** Comparison of overall scores for unsupervised methods between the baseline simulation (*n* = 120) and simulations with less samples (*n* = 30). **(B)** Comparison of overall scores for unsupervised methods between the baseline simulation (*α*_0_ = 10) and simulations with less (*α*_0_ = 30, left panel) or more (*α*_0_ = 3, right panel) dispersion. Methods for which the difference in scores is significant are flagged with stars indicating the p-value. **(C)** Overall scores of supervised methods in the case of a missing (“-1 type”) or extra cell type (“+1 type”) in the reference matrix versus the correct cell types (“All types”). The yellow DNA icon stands for methylation data, the blue single-strand one for transcriptome data.

Unsupervised methods are also sensitive to the dispersion parameter. Indeed, all methods performed worse when there was less dispersion (*α*_0_ = 30) than in the baseline, and it is particularly true for the methylation where the difference in scores is always significant (Figure 5B). In the *α*_0_ = 3 situation, many methods perform better than the baseline except for ICA, and debCAM for methylation data and CDSeq for transcriptomic data, although the deterioration of the score was never significant. This improvement of the performance makes sense as well, as more dispersion means more information (Supplementary Figure 8): in the extreme situation where the *α*_0_ factor is very large, this would cause all samples to have the same composition [19]. In the case where one expects its samples to have a similar composition, the results from our benchmark indicates that deconvolution is less affected with RNA than with DNAm data.

We evaluated the effect of removing or adding a cell type in the reference profile matrix during supervised deconvolution (see Methods). As expected, the optimal performance is met when the reference profiles comprise only the cell types present in the samples (Figure 5C). This is true for all methods and both omics. Moreover, we observed that in the case of an extra cell type not present in the samples, rankings are conserved compared to having the correct cell types in the reference, except for FARDEEP which ranks 2 positions higher in transcriptome deconvolution. In the case of a missing cell type, rankings are modified, although overall scores remain close. For example, RLR ranks third in methylation deconvolution but its overall score (OS = 0.557) is close to those of the first and second methods, respectively InstaPrism (OS = 0.570) and NNLS (OS = 0.569). Methods which are the more robust to an incorrect reference (defined as a difference of less than 0.1 in *OS*) are DeconRNASeq and InstaPrism (for RNA data) in the case of a missing cell type. In the case of an extra cell type, the most robust ones are DeconRNASeq, InstaPrism, WISP, both FARDEEP methods, NNLS, SVR (for RNA data) and RLR (for both omics). On the other hand, methods which are the most sensitive to an incorrect reference (defined as a difference of more than 0.25 in *OS*) are the methylation methods CIBERSORT and epidishCP.

### Rare cell types detection strongly depends on the deconvolution method

Depending on the research question, the main goal could be to estimate rare cell types. For example, it is important to quantify the immune infiltrate in the tumour microenvironment where these cells are usually lowly abundant. We defined as rare cell types in our simulations those with proportions *α*_*i*_ below 5% (Supplementary Table 3). It encompassed 4 datasets (3 simple-omic, 1 multi-omic). We investigated whether our methods were fit to detect rare occurrences based on cell-type Pearson correlations. Starting with methylation deconvolution, we observed that all methods were less efficient to estimate the proportions of rare cell types compared to common ones except for RLR in the LuCL dataset (Figure 6 upper panel). Apart from this particular case, there are only few methods for which the detection of rare cell types is not too degraded as compared to the detection of common cell types (defined by a difference in mean correlation below 0.2): RLR, NNLS and CIBERSORT which are only supervised methods. On the other hand, methods for which the detection of rare cell types is strongly degraded (defined by a difference in mean correlation above 0.4) include InstaPrism, epidishCP, ICA, NMF, and debCAM. Interestingly, the ability to detect rare types does not depend linearly on the overall ranking, nor on the rankings of each dataset (Supplementary Figure 6), but rather on the class of the method: supervised methods are more efficient than unsupervised ones to detect rare types. Our conclusions hold true when we only compared the efficacy to detect rare cell types: RLR, NNLS and CIBERSORT are the best methods independently of their relative performance on common cell types.

**Fig. 6.**
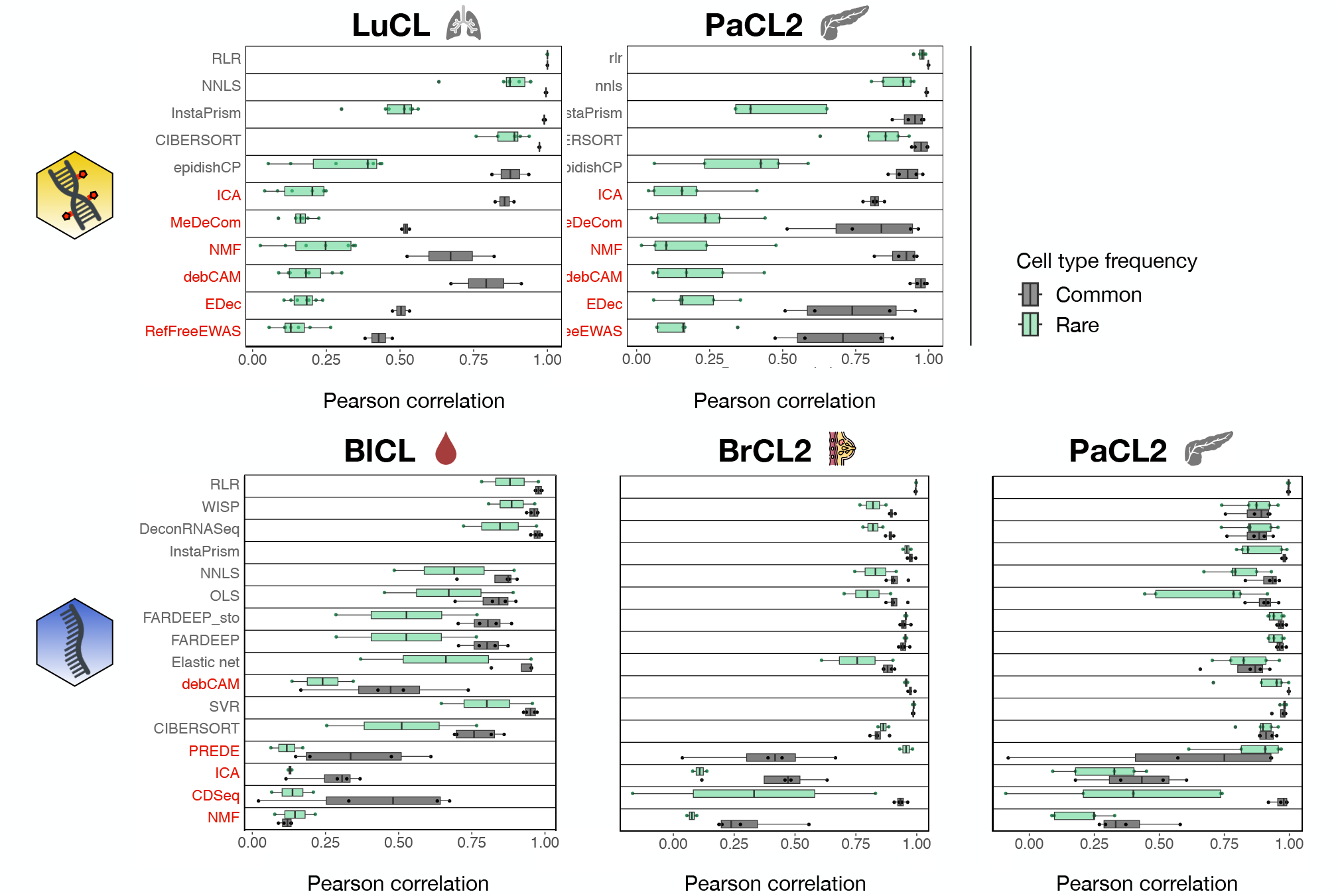
Investigation on the efficacy to detect rare cell type. Cell-type Pearson correlations for rare and common cell types were computed for each dataset containing rare cell types. Methods are ordered from top to bottom by their overall benchmark score for each omic type. Unsupervised methods are labelled in red. Number of rare types varies across datasets: n=7 for LuCL, n=5 for PaCL2, n=2 for BlCL and BrCL2. For the common types, n=2 for LuCL and n=4 for all other datasets. The yellow DNA icon stands for methylation data, the blue single-strand one for transcriptome data.

For RNA deconvolution, we also looked into performance degradation for the detection of rare cell types. With this omic type, the estimation of rare types is not systematically worse than the estimation of common types (Figure 6 lower panel), with some rare types being more correctly estimated than common types: e.g. SVR and PREDE in both cancer datasets. There is also less agreement across datasets than for the methylation. However, the mean correlation is in most cases smaller for rare types than for common types (74%, 35/47). For transcriptomic data, and since performances are better than for DNAm data, we defined reasonable performance degradation as a difference in mean correlation below 0.1. With this definition, 2 methods are not too degraded when detecting rare types: RLR and WISP. Conversely, and defining large degradation as a difference above 0.4, only CDSeq meets this criterion. On an absolute basis, RLR is the best method across datasets to estimate rare cell types. Finally, and based on the analysis of the multi-omic dataset PaCL2, we observed that rare cell types detection is the most competitive on RNA data. This relates to the fact that the transcriptome is already easier to deconvolve in general, as mentioned earlier.

### *In vitro* and *in vivo* datasets recapitulate *in silico* ranking

As mentioned earlier, simulations differ from real-life datasets and we need to ensure that our results are also valid for other data sources. We confronted in Figure 7 rankings obtained using datasets from different sources: gold-standard *in silico* and *in vitro* datasets, silver-standard *in vivo* datasets. We computed the ranking for each source independently of the others, as we do not want to compare the respective performance between sources but the ranking of the different methods itself. Of note, we cannot compute the secondary stability metrics for real datasets since we do not have replicates, so we have two different sets of metrics depending on the source (see Methods and Supplementary Figure 2). We also retained only the methods that ran on all *in vitro* and *in vivo* datasets, otherwise it would have favoured methods that were running only on few datasets (Supplementary Figure 9). Indeed, some methods were not able to deconvolve few real datasets, and we penalized it by excluding them in this comparison. We kept 9 out of the 11 DNAm methods and 12 out of the 16 RNA methods. InstaPrism was excluded in both omics as it requires to define variable cell types, which we chose to be the cancer cell types; hence it was run only on cancer datasets (see Methods).

**Fig. 7.**
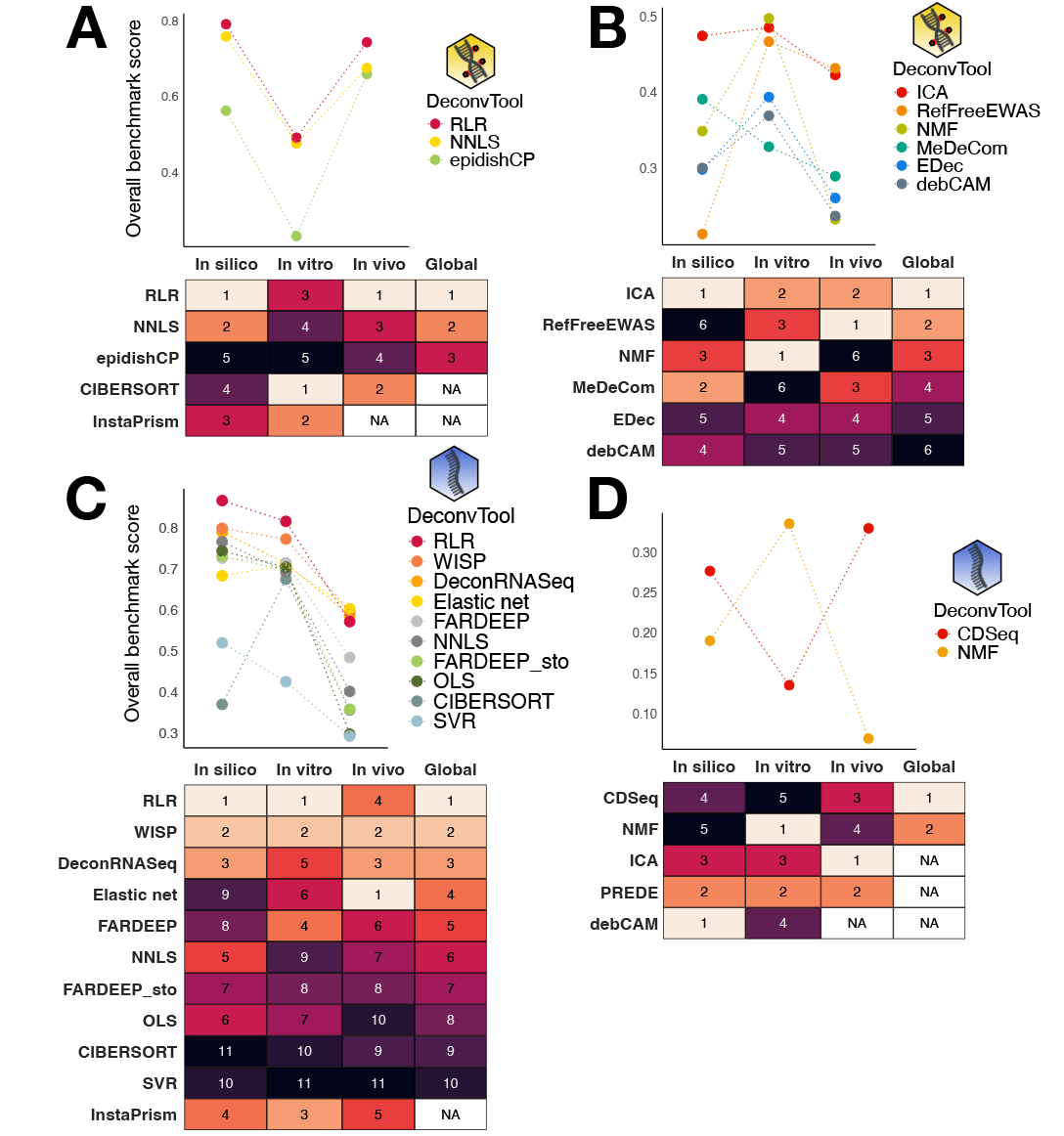
Rankings as a function of omic type, class and data source. Each panel displays the overall score of the methods in each source and the respective rankings, ‘global’ being the ranking as defined by the mean of the scaled overall scores for each method, for DNAm **(A**,**B)** and RNA **(C**,**D)** methods, supervised **(A**,**C)** or unsupervised **(B**,**D)**. The yellow DNA icon stands for methylation data, the blue single-strand one for transcriptome data. Methods that were excluded from the comparison across data sources or did not run on all datasets of a given source have no ranking (NA in the corresponding cell of the ranking tables)

For the supervised methylation methods, RLR remains the best method across all sources. For the unsupervised case, the best overall method is ICA, as outlined in the *in silico* ranking. Furthermore, we observed that the 3 best methods in each source are in the top 4 of the global ranking. Major differences are the rank change of RefFreeEWAS (6th in the *in silico* ranking), MeDeCom (6th in the *in vitro* ranking) and NMF (5th in the *in vivo* ranking).

In the supervised RNA deconvolution, we obtained a coherent ranking. SVR and CIBERSORT are consistently the worst methods, while RLR, WISP and DeconRNASeq are always in the top 4 for each source. In the unsupervised case, we only retained 2 out of the 5 methods, making it difficult to challenge any observation.

This confirms the robustness of our ranking method, as well as the hegemony of some methods over the others, such as RLR in the supervised class.

### Our benchmark provides a robust and comprehensive overview of a method’s performance

Thanks to our ranking process, we could unambiguously compare deconvolution tools. However, we can also look at intermediate and primary scores in order to better understand what makes a method efficient or not (Figure 8A,B,C,D, Supplementary Figure 10). For example, we observed that RLR performs well for all metrics in all except one *in vitro* DNAm dataset. We also observed that spiderplots of supervised methods looked more regular than for unsupervised ones, in terms of metrics and datasets. All those scores allowed us to come up with guidelines in terms of dataset’s easiness-of-deconvolution and algorithmic design’s strengths and weaknesses. It could also help choosing a method as a function of a specific use case. As an illustration, we compared InstaPrism and CIBERSORT to deconvolve PaCL2. InstaPrism made less errors (RMSE = 0.045, *R*^2^ = 0.93) than CIBERSORT (RMSE = 0.052, *R*^2^ = 0.92) on the whole matrix, and ran 10 times faster, but it failed to correctly estimate rare proportions compared to CIBERSORT, except for macrophages (Figure 8E,F,G). One should favour one method over the other as a function of its goal: if the goal is to predict the abundance of rare cell types and scalability is not an issue, the best tool is CIBERSORT. On the opposite, if the goal is to make less errors globally, InstaPrism is more efficient.

**Fig. 8.**
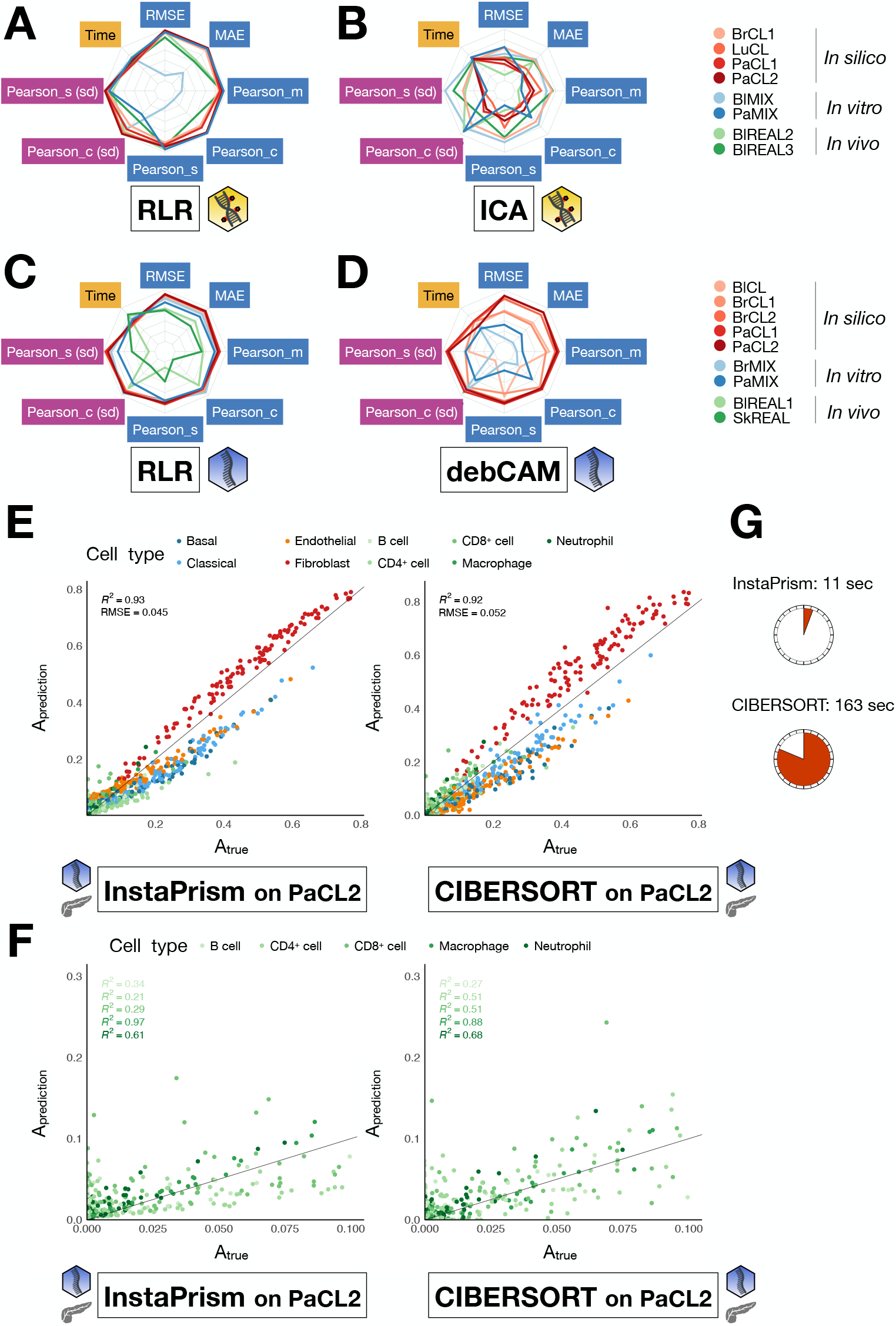
Detailed visualisation of methods’ performances. The benchmark allows to display several visual outputs. **(A**,**B**,**C**,**D)** Spiderplots of normalised-transformed primary metrics quantifying raw performance (blue), stability (violet) and scalability (yellow), for all dataset sources and for the two best methods in each class (supervised **(A**,**C)** and unsupervised **(B**,**D)**) and omic (methylation **(A**,**B)** and transcriptome **(C**,**D)**), as determined by the *in silico* ranking. **(E**,**F)** Predicted proportions as a function of true proportions for one simulation of PaCL2, coloured by cell type. All cell types are present in **(E)** and only rare types in **(F). (G)** Running time for each algorithm. Methods flagged with the yellow icon are run on the methylation, methods flagged with the blue icon on transcriptome.

## Methods

### Original PaCL2 and PaMIX datasets generation

This dataset consists of 30 mixtures and 9 pure cell types, constructed to recapitulate the heterogeneity seen in real pancreatic adenocarcinoma. The in vitro mixes contained variable proportions of (i) human tumour cells (CAPAN-1 and Mia PaCa-2), (ii) cancer associated fibroblasts (mix of 2 primary cell lines), (iii) human tumour derived-endothelial cells (HMEC) and immune cells that were FACS-sorted from healthy donors (B cells, CD4^+^ cells, CD8^+^ cells, neutrophils and M2-macrophages). All patients gave their written informed consent for the use of their specimen for research. Cells were mixed and RNA/DNA were simultaneously extracted followed by RNAseq (RNA-seq poly A) and methylome (MethEPIC 850K). For all samples, DNA/RNA were extracted using the ALLPrep tissue kit (Qiagen, Venlo, The Netherlands) following the manufacturer’s instructions.

RNA-seq libraries were prepared using the NEBNext Ultra II Directional RNA preparation kit, and paired-end 100-bp sequencing was conducted on the NovaSeq Illumina platform. Gene expression profiles were generated using Fastq files and aligned using STAR (2.7.1a) on UCSC hg38 genome. Bam files were counted using featureCounts (v2.0.0) with options -p -s 2 -T 15 -t exon -g gene_name.

Gene counts were normalized using standard DESeq2 procedure. DNA methylation was acquired according to standard Illumina protocol for Infinium Methylation EPIC BeadChip. The raw DNA methylation intensity data files (IDAT) were processed with the lumi and methylumi R packages. We performed pre-normalization filtering (removing probes containing SNP, high intensity probes and undetected probes) and normalization using colour balance adjustment and between-sample normalisation with the “quantile” method. The gene expression data and the DNA methylation data have been deposited on GEO under accession codes GSE281204 and GSE281305.

### Data simulation

We simulated expression/methylation bulk data 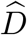 and proportion matrices and 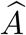, based on real reference profile matrices *T* and *a priori* knowledge on the proportions classically found in real samples (Supplementary Table 3). 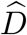 is of size *f × S, T* of size *f × K* and 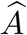 of size *K × S*, with *f* the number of features, *S* the number of samples and *K* the number of cell types.

We first generated proportions 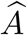 based on a Dirichlet distribution and *a priori* knowledge on realistic proportions(Supplementary Table 3):

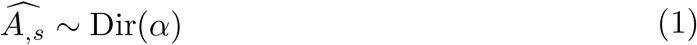

with 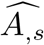 the vector of proportions of *K* cell types in the sample *s* and *α* = *α*_0_ *×* (*α*_1_, …, *α*_*K*_) the shape vector of the *K* cell types. For each dataset, *α* has been chosen proportional to prior knowledge on the cell types mix in real data, multiplied by a factor *α*_0_ controlling the dispersion around those typical proportions (Supplementary Table 3). The dispersion factor mimics biological noise: the higher the factor, the lower the dispersion around typical proportions.

Proportions 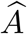 were then convoluted with reference profiles of pure cell types *T* to obtain the expression/methylation matrix 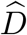. Table 2 indicates which references were chosen to simulate a given dataset. Finally we added a noise *ϵ* to model technical noise: a Negative Binomial noise for RNA and a Gaussian noise on M-values for DNAm data (see Supplementary Methods for the parameters of the Negative Binomial and Gaussian distributions) [17, 20, 30].

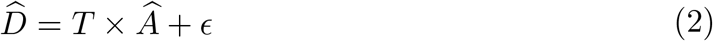

We generated sets of 10 replicates for each reference matrix *T* (Table 2).

Removal or addition of a cell type in the reference profile matrix were performed by modifying *T*, for the experiments of missing or extra cell types. To have a reference matrix with a missing cell type, we removed the first column of *T* ∈ ℝ^*f*×*K*^ after ordering the columns alphabetically. For the extra cell type, we generated a fake cell type based on the real ones. Basically, we selected all immune cell types, computed the mean profile and added noise (see Supplementary Methods).

### Computational deconvolution: formulation of the problem

The reference, proportions and expression/methylation matrices can be connected via the following equation:

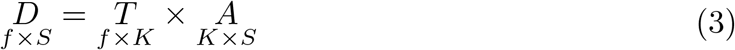

We can add further constraints of non-negativity (NN) and sum-to-one (STO) on

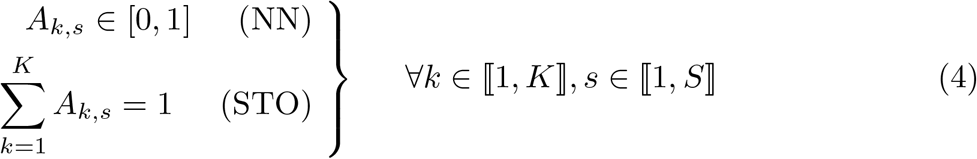

### Supervised deconvolution algorithms

We used multiple approaches for supervised deconvolution. Several methods are based on least squares (LS): Robust Linear Regression (RLR) [31], Non-Negative LS (NNLS) [32], epidishCP [31], Ordinary LS (OLS) [32], WISP [33], DeconRNASeq [34], Elastic net and FARDEEP [35]. We added an STO constraint on FARDEEP for the method FARDEEP sto. Other methods are based on a Support Vector Machine: CIBER-SORT [36] and Support Vector Regression (SVR) [32]. The last method relies on a Bayesian approach to directly sample the columns 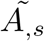 from the posterior distribu-tion *p*(*A*_,*s*_|*D*_,*s*_,*T*): InstaPrism [37]. InstaPrism requires to define variable cell types, which we chose to be the tumour types. Hence, InstaPrism was not run on non-cancer datasets.

For *in silico* data, we used the same reference matrix as the one employed for the simulations. For *in vitro* data, we used as reference the profiles of the pure cell types that were mixed. Except for BlMIX, the pure types were profiled at the same time as the mixes. For *in vivo* data, BlREAL1 and BlREAL2 were published along with matching references. We used the reference profiles from BlREAL2 to do supervised deconvolution of BlREAL3. For SkREAL, we used signatures accessed with the function *EPIC::TRef* (Table 2, column “Reference”).

### Unsupervised deconvolution algorithms

Unsupervised deconvolution was conducted using two primary approaches. In the first approach, there is no assumptions about the reference matrix, and both *A* and *T* are estimated simultaneously through Independent Component Analysis (ICA). In the second approach, *A* and *T* are estimated alternatively using LS methods, including Non-negative Matrix Factorization (NMF), RefFreeEWAS [38], MeDeCom [39], EDec [40], and PREDE [41]. Other approaches included convex analysis with debCAM [42] or Latent Dirichlet Allocation (LDA) with CDSeq [43]. Although debCAM was initially designed for RNA data, we also tested its performance on DNAm data. For all unsupervised methods, we used the true number of cell types as the number of unsupervised components in the algorithms.

After estimating 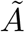, we identified the components by matching them to the cell types in the ground truth matrix *A*. Specifically, we aligned rows from 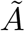 to those from the ground truth *A* by solving a linear sum assignment problem, using the R package *clue* to order rows in a way that maximizes correlations.

### Implementation of deconvolution methods: feature selection and execution of deconvolution algorithms

Feature selections were performed using the *TOAST* R package. The most variable features were identified with the function *findRefinx* and TOAST discriminating features with the function *csDeconv*, using ICA for the “FUN” argument.

Most algorithms were executed with default parameters (see Supplementary Methods for more details). The deconvolution analyses were run with a Snakemake workflow within an Apptainer container on the University of Grenoble computing infrastructure GRICAD (https://gricad.univ-grenoble-alpes.fr). Each analysis was run on a single CPU node with 32 cores allocated for parallelized methods. The seed was fixed for reproducibility. To accelerate computation, deconvolution for methylation data was limited to the 30,000 most variable features as captured by the function *TOAST::findRefinx*.

### Performance metrics

We calculated several metrics grouped in three categories, namely:

- Raw performance: RMSE, MAE, Pearson correlation *P* on the whole predicted matrix 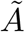, median across cell types of the Pearson correlations on cell types *P*_*k*_, median across samples of the Pearson correlations on samples *P*_*s*_;
- Stability across replicates: standard deviation of RMSE, MAE and *P*, median across replicates of the standard deviation of *P*_*k*_ and *P*_*s*_;
- Time: Per-sample time of deconvolution, log-transformed, and its standard deviation across replicates.

Secondary stability metrics are calculated exclusively on replicates. The ranking process that includes both primary and secondary metrics is referred to as ‘A,’ while the process based solely on primary metrics is referred to as ‘B’ (Supplementary Figure 2F). When replicates were available, primary metrics were averaged by taking the median across replicates (see Supplementary Methods).

We normalized the metrics as described in [44]. Let *M*^*X*^ be the method *×* metric matrix for a dataset *X*. Each method is a combination of a feature selection strategy and a deconvolution algorithm. We first normalized each metric for the dataset *X* for all methods such that the different scores are all between 0 and 1, and transformed it such that 1 is the best score and 0 the worst (see Supplementary Methods). After normalization-transformation, Pearson correlation metrics (resp. standard deviation of Pearson correlations) are merged into a Pearson meta-score (resp. Pearson stability meta-score) with the arithmetic mean (refer to Supplementary Methods). The output of this normalization - transformation - merging step is 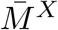.

### Ranking processes

We designed a consensus ranking processes to aggregate the different metrics into an overall score. The global aggregation process takes the method *×* metric matrices for all datasets as input, and outputs a vector of methods’ overall scores *S* via a three-steps process. First, metrics are aggregated using the geometric mean for each method, dataset and category. Second, we compute a weighted geometric mean to aggregate across categories for a given method and dataset, with weights of 1 for the raw performance and 0.5 for the scalability and stability. Finally, we aggregate across datasets via the arithmetic mean for each method (see Supplementary Methods).

We tested three different inputs for the global aggregation described above (Supplementary Figure 1). (1) For the process of successive aggregations *S*_*raw*_, the input is 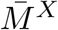. (2) For the process of average ranks *S*_*rank*_, the input is the matrix of ranks *R*^*X*^ computed separately for each metric from 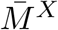 (Supplementary Methods and Supplementary Figure 1). (3) For the topsis process *S*_*topsis*_, the input is the matrix of TOPSIS scores computed from 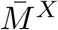. The TOPSIS score 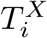 measures a ratio of the distances between each method *i* and archetypes of the best and worst possible methods for each dataset *X* (Supplementary Methods and Supplementary Figure 1) [25].

Finally, *S*_*consensus*_ is the arithmetic mean of the overall scores computed from each previously described processes *S*_*raw*_, *S*_*rank*_ and *S*_*topsis*_. We used the process *S*_*consensus*_ in all figures.

To test the quality of our ranking process, we quantified on *S*_*raw*_, *S*_*rank*_, *S*_*topsis*_ and *S*_*consensus*_ a series of criteria in order to characterize their behaviour [24]. We computed the average rank which is the normalized average rank of the winner, the Condorcet rate which is the rate of ranking the Condorcet winner first when one exists, and the generalization criterion which quantifies of how much a ranking depends on the set of metrics used (Supplementary Methods).

### Statistical significance of the overall score

We used a permutation test in order to associate p-values to the overall scores. For each pair of methods *i, j*, with *i* having a higher overall score than *j*, we did a one-sided test measuring how significant the difference in score was. We randomly shuffled their scores *M*_*i*,_ and *M*_*j*,_ 1,000 times. For each iteration *k*, we computed the overall scores 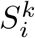 and 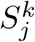 using the process *S*_*consensus*_, and the associated test statistic 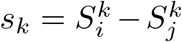. Finally, we compared the observed test statistic *s*_0_ = *S*_*i*_ *− S*_*j*_ to the distribution *{s*_*k*_,*k ∈* ⟦1, 1000⟧*}*.

## Code and data availability

The code for the deconvolution pipeline along with the Apptainer container definition file, and the code to reproduce the figures for the ranking pipeline are available on GitHub (https://github.com/bcm-uga/DeconvBenchmark) and Zenodo (DOI 10.5281/zenodo.14024479). The deconvolution pipeline can be adapted to add new methods and datasets. Similarly, the ranking pipeline can be modified. The accession codes for the datasets used in this study are listed in Table 2 and the directions on where to retrieve the proportion matrices for the *in vitro* mixes are given on our GitHub. The original dataset can be retrieved from GEO under the accession codes GSE281204 and GSE281305.

## Discussion

With this benchmark, we conducted a thorough evaluation of deconvolution algorithms across multiple omic types, addressing several critical aspects of their performance, such as robustness, scalability, and precision. The proposed framework evaluated both supervised and unsupervised approaches using diverse in silico, in vitro and in vivo benchmark datasets, including an original multi-omics dataset. The analysis spanned the impact of simulation parameters, the detection of rare cell types, and the consistency of algorithms ranking across diverse data sources. This comprehensive evaluation allowed us to tackle some of the major challenges in selecting deconvolution methods for transcriptomic and methylomic data, while paving the way for the development of new multi-omic algorithms, a significant step forward compared to existing benchmarks.

While our benchmark represents a notable advancement, our approach has certain limitations. The use of simulated data allowed us to work with large datasets with the exact ground truth, which is essential for rigorous evaluation. Although we applied common strategies for simulating technical noise, such as negative binomial noise for RNA data [17] and Gaussian noise for DNAm data [20, 30], we acknowledge that our simulations may not fully capture the complexity of real data. For instance, dependencies between genes and CpG probes or interactions between cells are not adequately modelled. These factors could partially explain the discrepancies in algorithms ranking across different datasets. Additionally, the scarcity of real-world datasets with reliable ground truth, and the uncertainty surrounding their accuracy, may also contribute to these inconsistencies [6].

Our benchmark allowed us to compare results with those from previous studies, revealing both similarities and differences. Consistent with our benchmark, [20] and [21] also observed that RLR was the best method for deconvolution of methylation data. However, our results diverged from other studies in some respects: for example, DeconRNASeq performed better in our evaluation than reported by [7] while it is the opposite for CIBERSORT (ranked last in our benchmark). These discrepancies likely stem from differences in the datasets used and the evaluation metrics chosen [6]. Our benchmark includes a larger variety of datasets and applies an overall ranking that merges multiple performance metrics, providing a more holistic view of algorithm performance.

The overall benchmark score allowed us to provide a single, interpretable ranking for each method, which simplifies comparison. However, this approach comes with trade-offs. Aggregating multiple metrics into a single score means that nuances from individual metrics are inevitably lost. To address this, we recommend that users refer back to the individual metrics in cases where a specific aspect of performance, *e*.*g*. speed or rare cell types detection, predominates. This flexibility allows for a more tailored selection of algorithms based on specific research needs. Additionally, our ranking method involved several subjective choices, such as selecting and categorizing metrics, along with weighting, and normalization procedure. While these choices were informed by existing benchmarks and the literature, they influence the final ranking. We chose metrics that were classically used in other benchmarks: RMSE and MAE measure the average error between the prediction and the reality, with RMSE increasing to a greater degree than MAE in case of a few large differences, and Pearson correlation measures the linear correlation. Some benchmarks also used Jensen-Shannon divergence [45], Aitchison distance [46] or sMAPE [16, 21]. Therefore, future studies could explore alternative ranking methodologies or adjust the weights to emphasize specific performance aspects depending on the analysis context. The flexibility of our benchmark ensures that as new methods and metrics emerge, it can be readily adapted to meet evolving research needs.

Our benchmark provides valuable guidance for researchers looking to select appropriate deconvolution methods based on their datasets. Datasets can be described by several characteristics: number of features, number of cell types, technology, etc (Supplementary Figure 11). We investigated if some characteristics could be linked to the easiness-of-deconvolution. We explored the sensitivity of each method to the datasets and observed that unsupervised methods tend to be more sensitive to the datasets than supervised ones (Supplementary Figure 12). Although our attempts to link specific dataset characteristics, such as heterogeneity, to deconvolution difficulty were inconclusive (Supplementary Figure 13), this remains a promising area for future research. Expanding the benchmark to include more datasets could help clarify these relationships.

Beyond its immediate application, our benchmark offers a rigorous framework that can be used to test and validate new deconvolution methods. Thanks to our strategy of giving a single rating per method we could unambiguously evaluate 20 algorithms across classes and omics. One of the key strengths of our benchmark is the inclusion of a dual-omic evaluation, which offers a unique perspective on how deconvolution methods perform across both transcriptomic and methylomic data. We have also observed that a consensus strategy did not outperform the best methods in our evaluation (Supplementary Figure 14), but this remains an area where future innovations could have a significant impact. By providing a reproducible and transparent framework, our benchmark can be reused and extended as new methods are developed, contributing to the continuous improvement of deconvolution techniques. Indeed, our benchmark is well-positioned to accommodate new types of data, such as spatial transcriptomics, which represent a growing frontier in the analysis of tumour heterogeneity.

Finally, our benchmark serves two key purposes: to evaluate the performance of existing deconvolution methods across a wide range of conditions; and to provide aflexible, extensible platform for the integration of future developments in the field. This evaluation was done on multiple datasets, including an original multi-omics dataset. By comparing algorithms across transcriptomic and methylomic data, we have laid the groundwork for the next generation of multi-omic deconvolution tools. Moreover, the ability to integrate additional datasets and algorithms ensures that this benchmark will remain a valuable resource in the future. Looking ahead, we anticipate that our framework could be extended to include spatial deconvolution methods, further broadening its applicability and relevance in the study of complex biological systems.

## Acknowledgements

We thank the GRICAD UGA mesocenter for computing resources. We thank Florent Petitprez for his insights into the design of the in vitro mixture dataset.

## Contributions

Conceptualization and methodology: E.A., Y.B. and M.R. Data generation and curation: F.C., M.A., A.B., L.A., Y.K and J.C. Project administration: Y.B. and M.R. Data analysis: E.A., V.B., L.M.P. and S.K. Writing: E.A., Y.B. and M.R. All authors have reviewed the manuscript and agreed to its publication.

## Competing interests

Authors declare that they have no competing interests.

## Funding

This work is a contribution of the EIT Health program COMETH. It has been partially supported by MIAI @ Grenoble Alpes (ANR-19-P3IA-0003), and by the French Agency for National Research (CauseHet // ANR-22-CE45-0030). Finally, it has also been carried out with financial support from ITMO Cancer of Aviesan within the framework of the 2021-2030 Cancer Control Strategy, on funds administered by Inserm (ACACIA project AAP-MIC-2021).

## Notes

https://github.com/bcm-uga/DeconvBenchmark

### Competing Interest Statement

The authors have declared no competing interest.

## References

[1] Shen-Orr, S.S., Gaujoux, R.: Computational deconvolution: extracting cell typespecific information from heterogeneous samples. Current Opinion in Immunology 25(5), 571–578 (2013) 10.1016/j.coi.2013.09.015

[2] Finotello, F., Trajanoski, Z.: Quantifying tumor-infiltrating immune cells from transcriptomics data. Cancer immunology, immunotherapy: CII 67(7), 1031–1040 (2018) 10.1007/s00262-018-2150-z

[3] De Visser, K.E., Joyce, J.A.: The evolving tumor microenvironment: From cancer initiation to metastatic outgrowth. Cancer Cell 41(3), 374–403 (2023) 10.1016/j.ccell.2023.02.016

[4] Schwartz, R., Shackney, S.E.: Applying unmixing to gene expression data for tumor phylogeny inference. BMC Bioinformatics 11(1), 42 (2010) 10.1186/1471-2105-11-42

[5] Zheng, S.: Benchmarking: contexts and details matter. Genome Biology 18(1), 129 (2017) 10.1186/s13059-017-1258-3

[6] Garmire, L.X., Li, Y., Huang, Q., Xu, C., Teichmann, S.A., Kaminski, N., Pellegrini, M., Nguyen, Q., Teschendorff, A.E.: Challenges and perspectives in computational deconvolution of genomics data. Nature Methods 21(3), 391–400 (2024) 10.1038/s41592-023-02166-6

[7] Avila Cobos, F., Alquicira-Hernandez, J., Powell, J.E., Mestdagh, P., De Preter, K.: Benchmarking of cell type deconvolution pipelines for transcriptomics data. Nature Communications 11(1), 5650 (2020) 10.1038/s41467-020-19015-1

[8] Nadel, B.B., Oliva, M., Shou, B.L., Mitchell, K., Ma, F., Montoya, D.J., Mouton, A., Kim-Hellmuth, S., Stranger, B.E., Pellegrini, M., Mangul, S.: Systematic evaluation of transcriptomics-based deconvolution methods and references using thousands of clinical samples. Briefings in Bioinformatics 22(6), 265 (2021) 10.1093/bib/bbab265

[9] Vallania, F., Tam, A., Lofgren, S., Schaffert, S., Azad, T.D., Bongen, E., Haynes, W., Alsup, M., Alonso, M., Davis, M., Engleman, E., Khatri, P.: Leveraging heterogeneity across multiple datasets increases cell-mixture deconvolution accuracy and reduces biological and technical biases. Nature Communications 9(1), 4735 (2018) 10.1038/s41467-018-07242-6

[10] He, D., Chen, M., Wang, W., Song, C., Qin, Y.: Deconvolution of tumor composition using partially available DNA methylation data. BMC Bioinformatics 23(1), 355 (2022) 10.1186/s12859-022-04893-7

[11] Houseman, E.A., Kile, M.L., Christiani, D.C., Ince, T.A., Kelsey, K.T., Marsit, C.J.: Reference-free deconvolution of DNA methylation data and mediation by cell composition effects. BMC Bioinformatics 17(1), 259 (2016) 10.1186/s12859-016-1140-4

[12] Yamawaki, T.M., Lu, D.R., Ellwanger, D.C., Bhatt, D., Manzanillo, P., Arias, V., Zhou, H., Yoon, O.K., Homann, O., Wang, S., Li, C.-M.: Systematic comparison of high-throughput single-cell RNA-seq methods for immune cell profiling. BMC Genomics 22(1), 66 (2021) 10.1186/s12864-020-07358-4

[13] Hippen, A.A., Omran, D.K., Weber, L.M., Jung, E., Drapkin, R., Doherty, J.A., Hicks, S.C., Greene, C.S.: Performance of computational algorithms to deconvolve heterogeneous bulk ovarian tumor tissue depends on experimental factors. Genome Biology 24(1), 239 (2023) 10.1186/s13059-023-03077-7

[14] Avila Cobos, F.A., Panah, M.J.N., Epps, J., Long, X., Man, T.-K., Chiu, H.-S., Chomsky, E., Kiner, E., Krueger, M.J., Bernardo, D., Voloch, L., Mole-naar, J., Hooff, S.R., Westermann, F., Jansky, S., Redell, M.L., Mestdagh, P., Sumazin, P.: Effective methods for bulk RNA-seq deconvolution using scnRNA-seq transcriptomes. Genome Biology 24(1), 177 (2023) 10.1186/s13059-023-03016-6

[15] Dietrich, A., Merotto, L., Pelz, K., Eder, B., Zackl, C., Reinisch, K., Edenhofer, F., Marini, F., Sturm, G., List, M., Finotello, F.: Benchmarking second-generation methods for cell-type deconvolution of transcriptomic data. “Preprint at https://www.biorxiv.org/content/10.1101/2024.06.10.598226v1” (2024)

[16] Zhang, W., Zhang, X., Liu, Q., Wei, L., Qiao, X., Gao, R., Liu, Z., Wang, X.: Deconer: A comprehensive and systematic evaluation toolkit for reference-based cell type deconvolution algorithms using gene expression data. “Preprint at https://www.biorxiv.org/content/10.1101/2023.12.24.573278v1” (2023)

[17] Jin, H., Liu, Z.: A benchmark for RNA-seq deconvolution analysis under dynamic testing environments. Genome Biology 22(1), 102 (2021) 10.1186/s13059-021-02290-6

[18] Sturm, G., Finotello, F., Petitprez, F., Zhang, J.D., Baumbach, J., Fridman, W.H., List, M., Aneichyk, T.: Comprehensive evaluation of transcriptome-based cell-type quantification methods for immuno-oncology. Bioinformatics 35(14), 436–445 (2019) 10.1093/bioinformatics/btz363

[19] Decamps, C., Privé, F., Bacher, R., Jost, D., Waguet, A., HADACA consortium, Houseman, E.A., Lurie, E., Lutsik, P., Milosavljevic, A., Scherer, M., Blum, M.G.B., Richard, M.: Guidelines for cell-type heterogeneity quantification based on a comparative analysis of reference-free DNA methylation deconvolution software. BMC bioinformatics 21(1), 16 (2020) 10.1186/s12859-019-3307-2

[20] Teschendorff, A.E., Breeze, C.E., Zheng, S.C., Beck, S.: A comparison of reference-based algorithms for correcting cell-type heterogeneity in Epigenome-Wide Association Studies. BMC bioinformatics 18(1), 105 (2017) 10.1186/s12859-017-1511-5

[21] Song, J., Kuan, P.-F.: A systematic assessment of cell type deconvolution algorithms for DNA methylation data. Briefings in Bioinformatics 23(6), 449 (2022) 10.1093/bib/bbac449

[22] Mölder, F., Jablonski, K.P., Letcher, B., Hall, M.B., Tomkins-Tinch, C.H., Sochat, V., Forster, J., Lee, S., Twardziok, S.O., Kanitz, A., Wilm, A., Holtgrewe, M., Rahmann, S., Nahnsen, S., Köster, J.: Sustainable data analysis with Snakemake. F1000Research (2021). 10.12688/f1000research.29032.2

[23] Singularity Developers (2021) Singularity. 10.5281/zenodo.1310023

[24] Pavao, A., Vaccaro, M., Guyon, I.: Judging competitions and benchmarks: a candidate election approach. Paper presented at ESANN 2021 (2021). 10.14428/esann/2021.ES2021-122

[25] Hwang, C.-L., Yoon, K.: Methods for Multiple Attribute Decision Making. In: Multiple Attribute Decision Making. Lecture Notes in Economics and Mathematical Systems, vol. 186, pp. 58–191. Springer, Berlin, Heidelberg (1981). 10.1007/978-3-642-48318-93

[26] Li, Z., Wu, H.: TOAST: improving reference-free cell composition estimation by cross-cell type differential analysis. Genome Biology 20(1), 1–17 (2019) 10.1186/s13059-019-1778-0

[27] Cai, M., Yue, M., Chen, T., Liu, J., Forno, E., Lu, X., Billiar, T., Celedón, J., McKennan, C., Chen, W., Wang, J.: Robust and accurate estimation of cellular fraction from tissue omics data via ensemble deconvolution. Bioinformatics 38(11), 3004–3010 (2022) 10.1093/bioinformatics/btac279

[28] Avila Cobos, F., Vandesompele, J., Mestdagh, P., De Preter, K.: Computational deconvolution of transcriptomics data from mixed cell populations. Bioinformatics 34(11), 1969–1979 (2018) 10.1093/bioinformatics/bty019

[29] Sutton, G.J., Poppe, D., Simmons, R.K., Walsh, K., Nawaz, U., Lister, R., Gagnon-Bartsch, J.A., Voineagu, I.: Comprehensive evaluation of deconvolution methods for human brain gene expression. Nature Communications 13(1), 1358 (2022) 10.1038/s41467-022-28655-4

[30] Chalise, P., Raghavan, R., Fridley, B.L.: InterSIM: Simulation tool for multiple integrative “omic datasets”. Computer Methods and Programs in Biomedicine 128, 69–74 (2016) 10.1016/j.cmpb.2016.02.011

[31] Teschendorff, A.E., Zheng, S.C.: EpiDISH. Bioconductor. (2017). 10.18129/B9.bioc.EpiDISH

[32] Pfister, S., Kuettel, V., Ferrero, E.: Granulator: Rapid Benchmarking of Methods for in Silico Deconvolution of Bulk RNA-seq Data. (2022). 10.18129/B9.bioc.granulator

[33] Blum, Y., Meiller, C., Quetel, L., Elarouci, N., Ayadi, M., Tashtanbaeva, D., Armenoult, L., Montagne, F., Tranchant, R., Renier, A., Koning, L., Copin, M.-C., Hofman, P., Hofman, V., Porte, H., Le Pimpec-Barthes, F., Zucman-Rossi, J., Jaurand, M.-C., Reyniés, A., Jean, D.: Dissecting heterogeneity in malignant pleural mesothelioma through histo-molecular gradients for clinical applications. Nature Communications 10(1), 1333 (2019) 10.1038/s41467-019-09307-6

[34] Gong, T., Szustakowski, J.D.: DeconRNASeq: a statistical framework for deconvolution of heterogeneous tissue samples based on mRNA-Seq data. Bioinformatics 29(8), 1083–1085 (2013) 10.1093/bioinformatics/btt090

[35] Hao, Y., Yan, M., Heath, B.R., Lei, Y.L., Xie, Y.: Fast and robust deconvolution of tumor infiltrating lymphocyte from expression profiles using least trimmed squares. PLOS Computational Biology 15(5), 1006976 (2019) 10.1371/journal.pcbi.1006976

[36] Newman, A.M., Liu, C.L., Green, M.R., Gentles, A.J., Feng, W., Xu, Y., Hoang, C.D., Diehn, M., Alizadeh, A.A.: Robust enumeration of cell subsets from tissue expression profiles. Nature Methods 12(5), 453–457 (2015) 10.1038/nmeth.3337

[37] Hu, M., Chikina, M.: InstaPrism: an R package for fast implementation of BayesPrism. “Preprint at https://www.biorxiv.org/content/10.1101/2023.03.07.531579v2” (2023)

[38] Houseman, E.A., Molitor, J., Marsit, C.J.: Reference-free cell mixture adjustments in analysis of DNA methylation data. Bioinformatics 30(10), 1431–1439 (2014) 10.1093/bioinformatics/btu029

[39] Lutsik, P., Slawski, M., Gasparoni, G., Vedeneev, N., Hein, M., Walter, J.: MeDeCom: discovery and quantification of latent components of heterogeneous methylomes. Genome Biology 18(1), 55 (2017) 10.1186/s13059-017-1182-6

[40] Onuchic, V., Hartmaier, R.J., Boone, D.N., Samuels, M.L., Patel, R.Y., White, W.M., Garovic, V.D., Oesterreich, S., Roth, M.E., Lee, A.V., Milosavljevic, A.: Epigenomic Deconvolution of Breast Tumors Reveals Metabolic Coupling between Constituent Cell Types. Cell Reports 17(8), 2075–2086 (2016) 10.1016/j.celrep.2016.10.057

[41] Qin, Y., Zhang, W., Sun, X., Nan, S., Wei, N., Wu, H.-J., Zheng, X.: Deconvolution of heterogeneous tumor samples using partial reference signals. PLOS Computational Biology 16(11), 1008452 (2020) 10.1371/journal.pcbi.1008452

[42] Wang, N., Hoffman, E.P., Chen, L., Chen, L., Zhang, Z., Liu, C., Yu, G., Herrington, D.M., Clarke, R., Wang, Y.: Mathematical modelling of transcriptional heterogeneity identifies novel markers and subpopulations in complex tissues. Scientific Reports 6(1), 18909 (2016) 10.1038/srep18909

[43] Kang, K., Meng, Q., Shats, I., Umbach, D.M., Li, M., Li, Y., Li, X., Li, L.: CDSeq: A novel complete deconvolution method for dissecting heterogeneous samples using gene expression data. PLOS Computational Biology 15(12), 1007510 (2019) 10.1371/journal.pcbi.1007510

[44] Saelens, W., Cannoodt, R., Todorov, H., Saeys, Y.: A comparison of single-cell trajectory inference methods. Nature Biotechnology 37(5), 547–554 (2019) 10.1038/s41587-019-0071-9

[45] Sang-aram, C., Browaeys, R., Seurinck, R., Saeys, Y.: Spotless: a reproducible pipeline for benchmarking cell type deconvolution in spatial transcriptomics. eLife 12 (2023) 10.7554/eLife.88431.1

[46] Tran, K.A., Addala, V., Johnston, R.L., Lovell, D., Bradley, A., Koufariotis, L.T., Wood, S., Wu, S.Z., Roden, D., Al-Eryani, G., Swarbrick, A., Williams, E.D., Pearson, J.V., Kondrashova, O., Waddell, N.: Performance of tumour microenvironment deconvolution methods in breast cancer using single-cell simulated bulk mixtures. Nature Communications 14(1), 5758 (2023) 10.1038/s41467-023-41385-5

[47] Azizi, E., Carr, A.J., Plitas, G., Cornish, A.E., Konopacki, C., Prabhakaran, S., Nainys, J., Wu, K., Kiseliovas, V., Setty, M., Choi, K., Fromme, R.M., Dao, P., McKenney, P.T., Wasti, R.C., Kadaveru, K., Mazutis, L., Rudensky, A.Y., Pe’er, D.: Single-Cell Map of Diverse Immune Phenotypes in the Breast Tumor Microenvironment. Cell 174(5), 1293–130836 (2018) 10.1016/j.cell.2018.05.060

