## Supplementary Figures and Tables for "A robust workflow to benchmark deconvolution of multi-omic data"

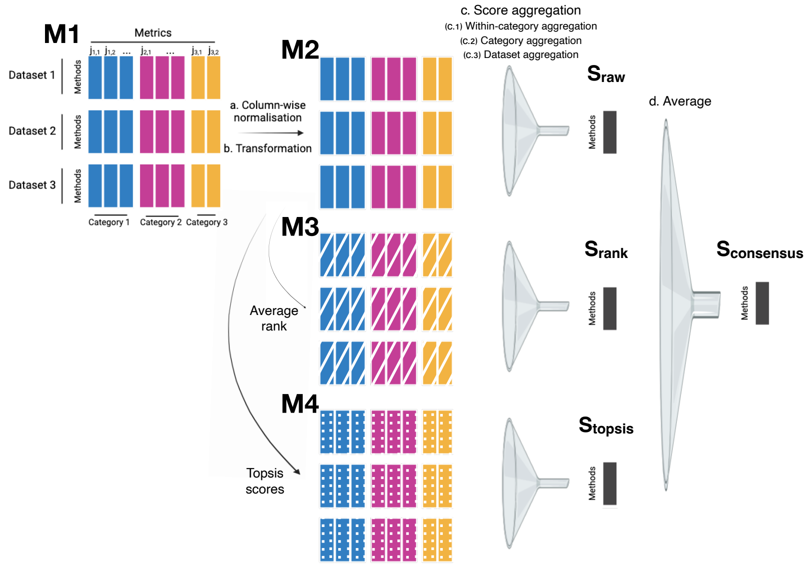


**Supplementary Figure 1** Design of the different ranking processes. The first step of normalisation (a) and transformation (b) on the metric-by-dataset-by-method score matrix **M1** is common to all processes. The matrix **M2** then goes through the global 3-steps aggregation process (c) described in Figure **??**C ($S_{raw}$), or is used to compute average ranks **(M3)** and topsis scores **(M4)**. Matrices **M3** and **M4** then go through the global aggregation process (c) (respectively $S_{rank}$ and $S_{topsis}$). The 3 ranking processes $S_{raw}$, $S_{rank}$ and $S_{topsis}$ are finally averaged (d) into $S_{consensus}$.

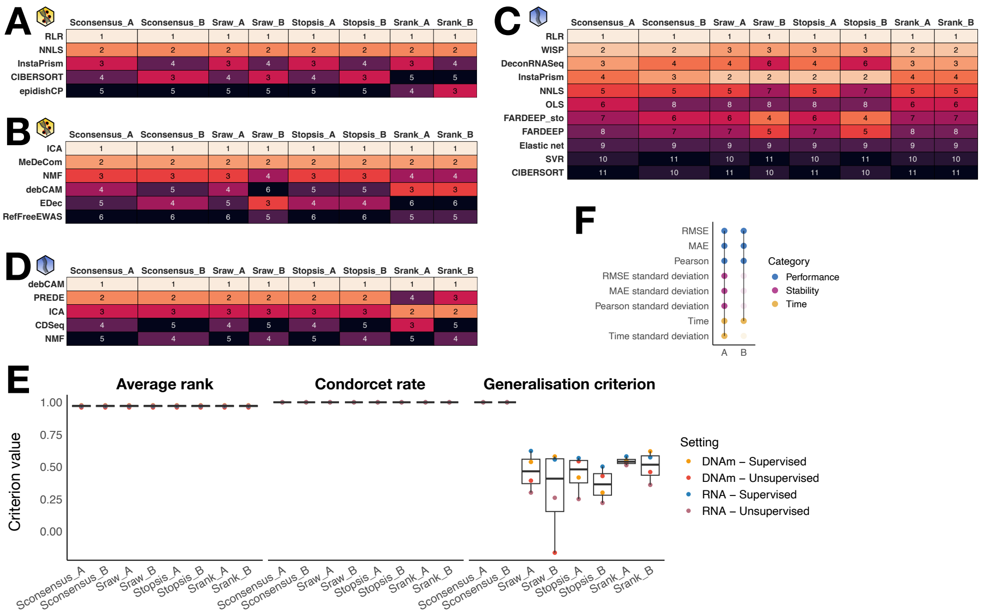


**Supplementary Figure 2** Comparison of the different ranking processes. Various ranking processes yield comparable rankings, especially for the top performing candidates. Ranking tables for the 4 ranking processes tested, for the methylation (**(A,B), yellow DNA icon** and transcriptome (**(C,D), blue single-strand icon** blocks and the supervised **(A,C)** or unsupervised **(B,D)** classes of methods. **(E)** Goodness-of-ranking criteria display similar scores for the different ranking processes, except for the generalisation criterion which shows the superiority of the consensus process. **(F)** Combinations A and B include different metrics.

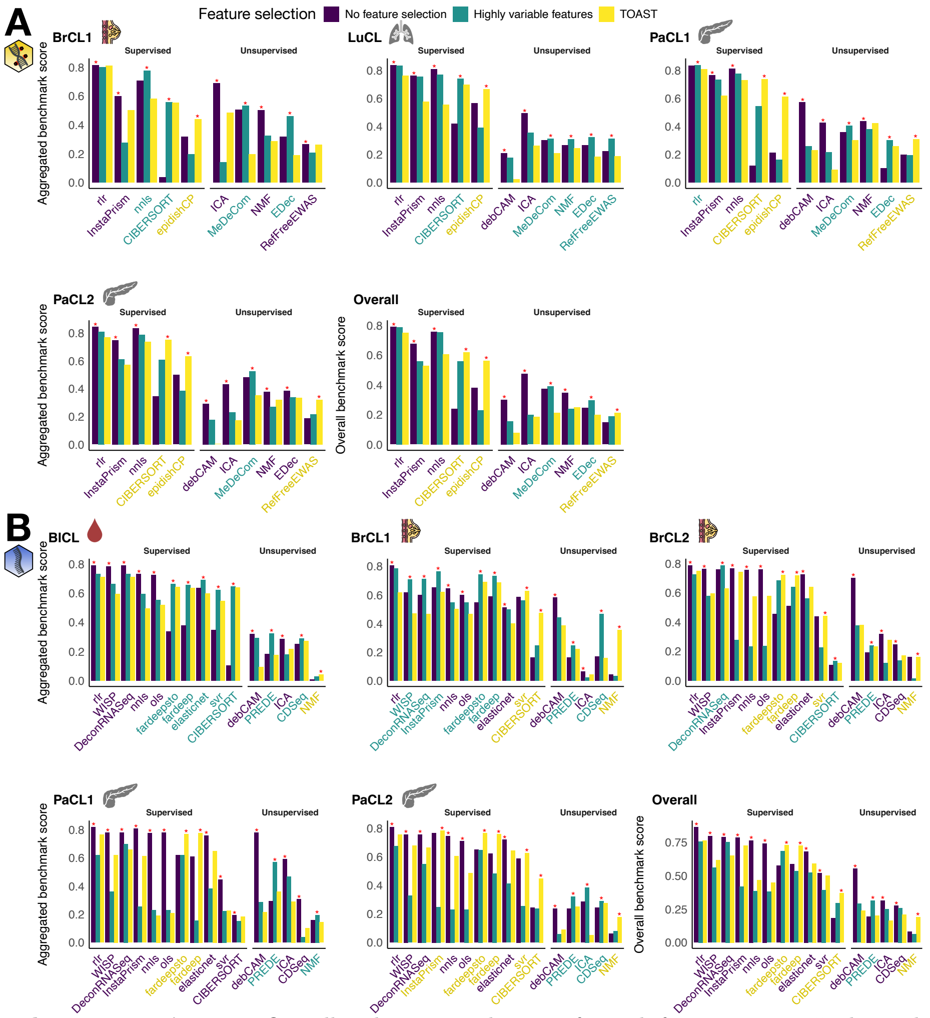


**Supplementary Figure 3** Overall and aggregated scores, for each feature setting. It shows that the no feature selection strategy performs the best in most cases, both for methylation **(A)**, flagged with the yellow DNA icon, and transcriptome **(B)** data, flagged with the blue single-strand one.

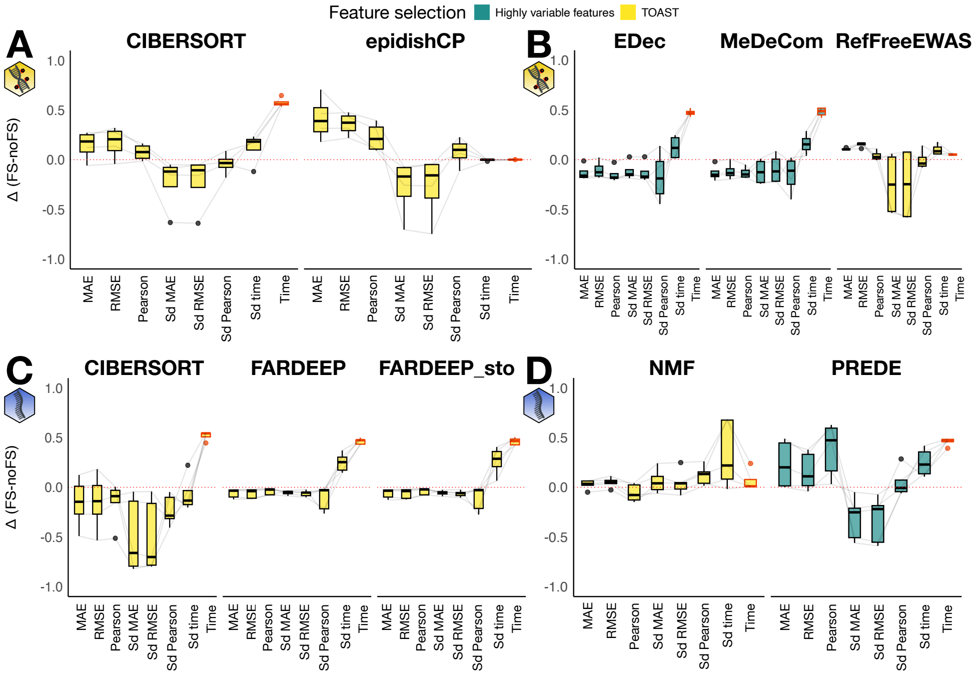


**Supplementary Figure 4** Detailed performance comparison with and without feature selection. Difference in normalised-transformed atomic scores between the best feature selection and the no feature selection for candidates for which feature selection improves the overall benchmark score in simulated data (refer to Supplementary Figure 3). Each data point is the difference in atomic score for a given dataset. The left panels **(A,C)** show supervised methods, the right ones **(B,D)** unsupervised methods. The upper panels **(A,B)** show the results for DNAm data (yellow DNA icon), the lower ones **(C,D)** for RNA data (blue single-strand icon). The time score is in red.

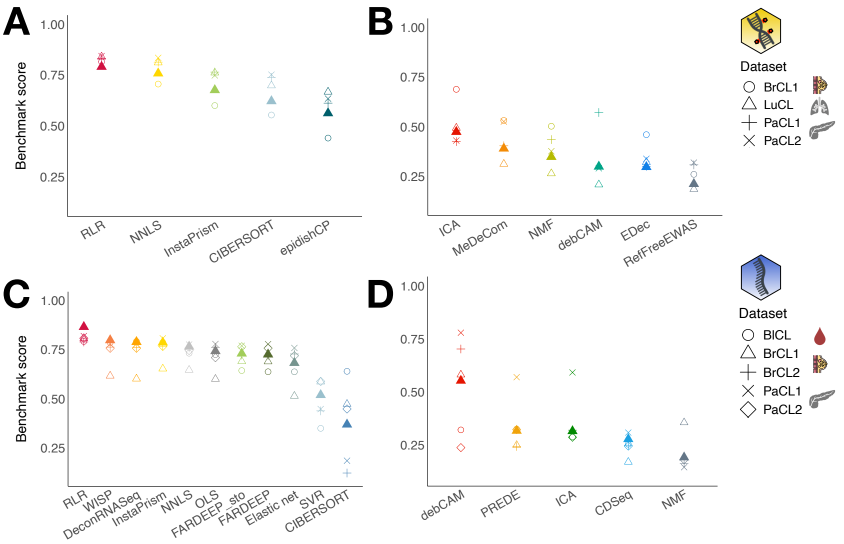


**Supplementary Figure 5** Overall and aggregated benchmark scores. Scores are for the methylation **(A,B)** and transcriptome **(C,D)** blocks in the supervised **(A,C)** or unsupervised **(B,D)** classes. The overall score is depicted by the full triangle symbol.

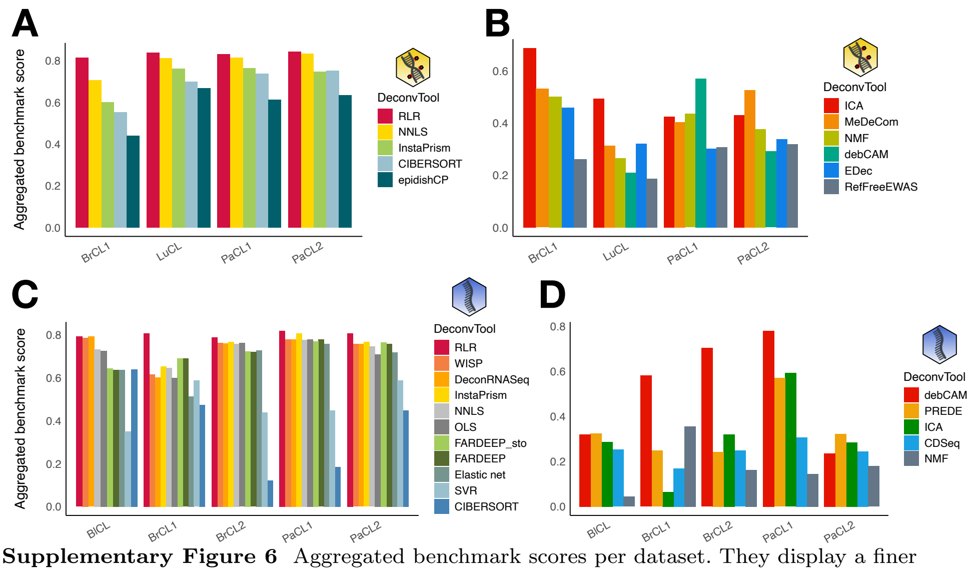


**Supplementary Figure 6** Aggregated benchmark scores per dataset. They display a finer granularity in the ranking of the different candidates, as a function of the omic (methylation **(A,B)**, transcriptome **(C,D)**) and class of methods (supervised **(A,C)**, unsupervised **(B,D)**).

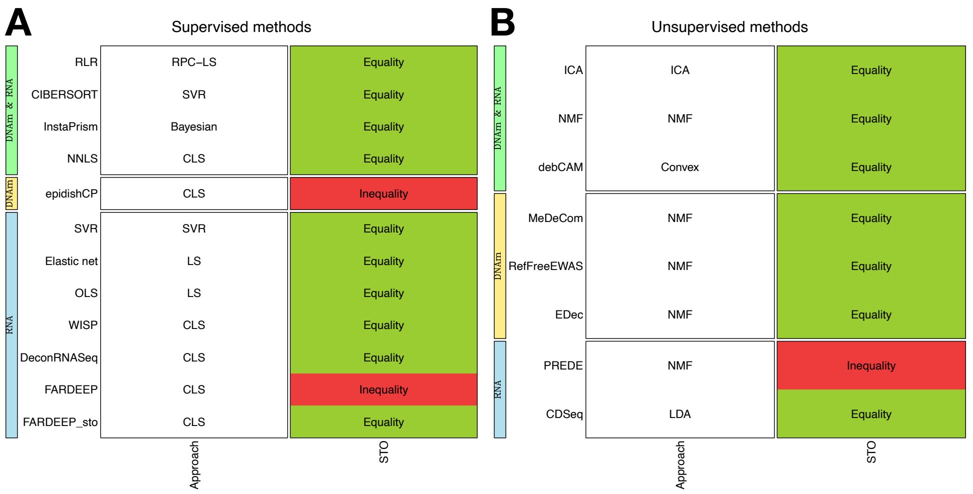


**Supplementary Figure 7** Different algorithmic designs exist. Designs for the supervised **(A)** and unsupervised **(B)** classes of methods. The second column indicates whether the sum-to-one (STO) constraint is a strict equality or an inequality.

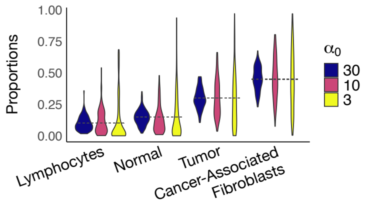


**Supplementary Figure 8** The dispersion factor affects the proportions' distribution. Different values of the dispersion factor $\alpha_0$ implies different ranges of variation around the proportions $\alpha_i, i \in [1, K]$ (dashed line) in the simulations. Example with the BrCL1 dataset.

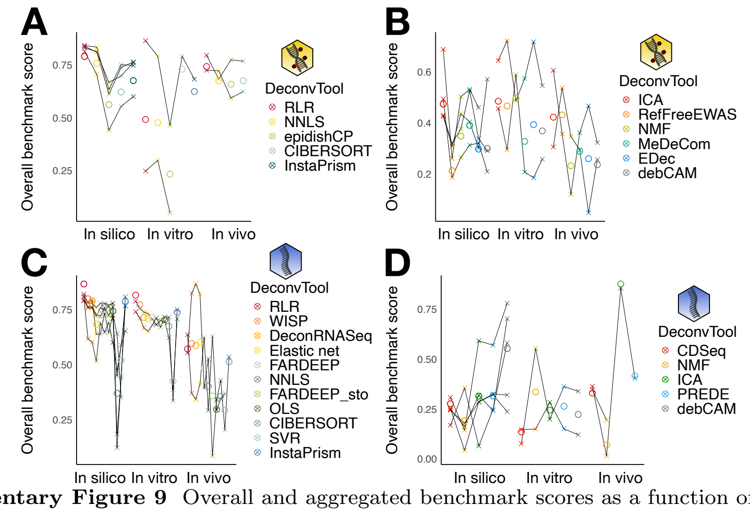


**Supplementary Figure 9** Overall and aggregated benchmark scores as a function of the data source. Overall scores are circles, aggregated scores are crosses, and datasets are linked by a full line, showing that some datasets can not be deconvolved by all tools. The left panels **(A,C)** display supervised methods, the right ones **(B,D)** unsupervised methods. The upper panels **(A,B)** display DNAm data, the lower ones **(C,D)** RNA data as depicted by the blue and yellow icons.

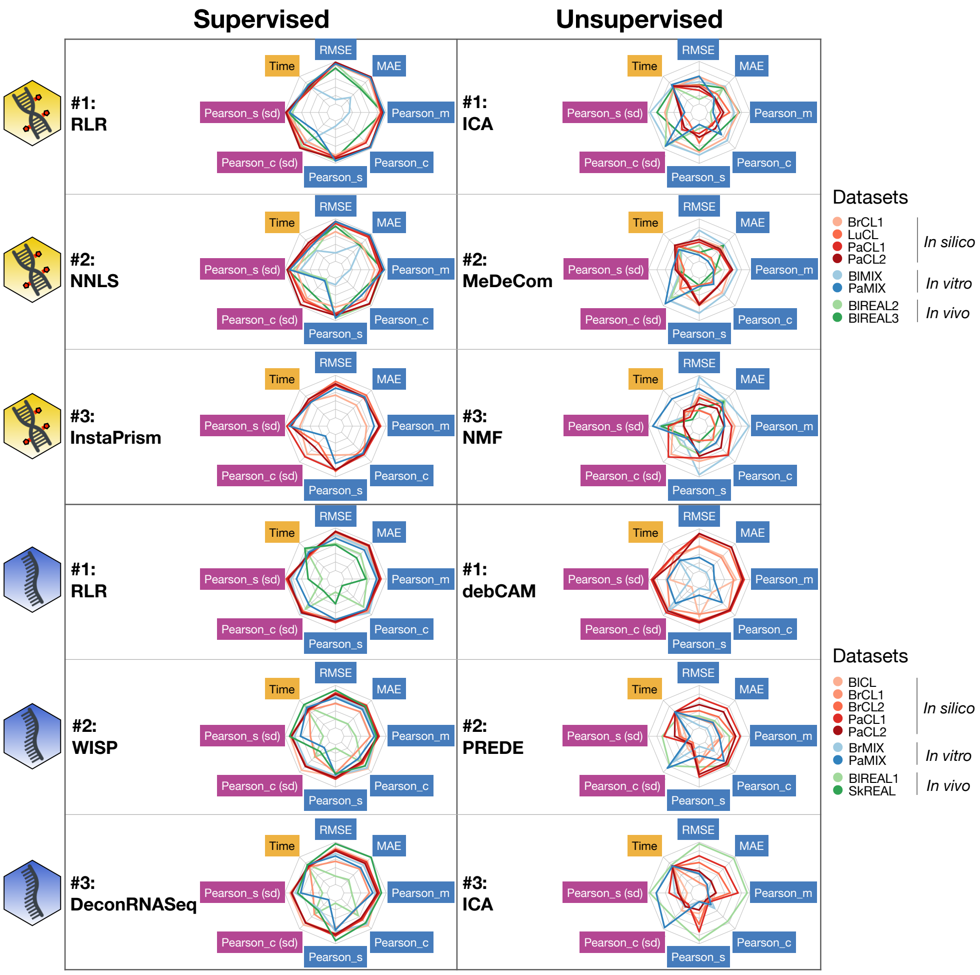


**Supplementary Figure 10** Spiderplots of the top 3 best methods in each setting. Methylation is flagged with the yellow DNA icon and transcriptome with the blue single-strand icon.

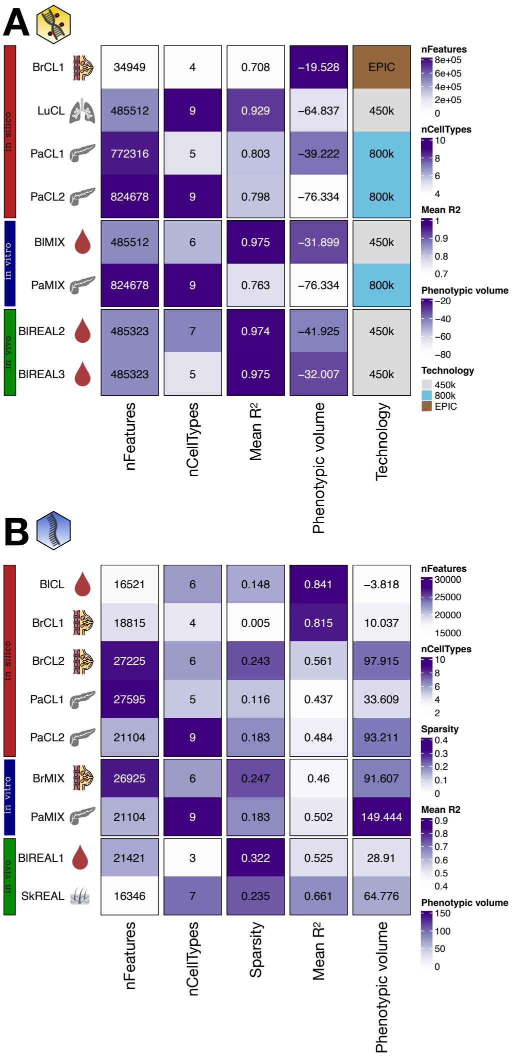


**Supplementary Figure 11** Datasets characteristics. (**A**) shows DNAm datasets and (**B**) RNA datasets. We computed for both omics the number of molecular features, the number of cell types, the mean Pearson correlation between cell types profiles, and the phenotypic volume as defined in [**?** ] (see Methods), the two latter serving as proxies of dataset heterogeneity. We also took into account the technology for the methylation block, while all transcriptomes were sequenced with the Illumina platform. For the transcriptome block, we added the sparsity, defined as the mean of null counts.

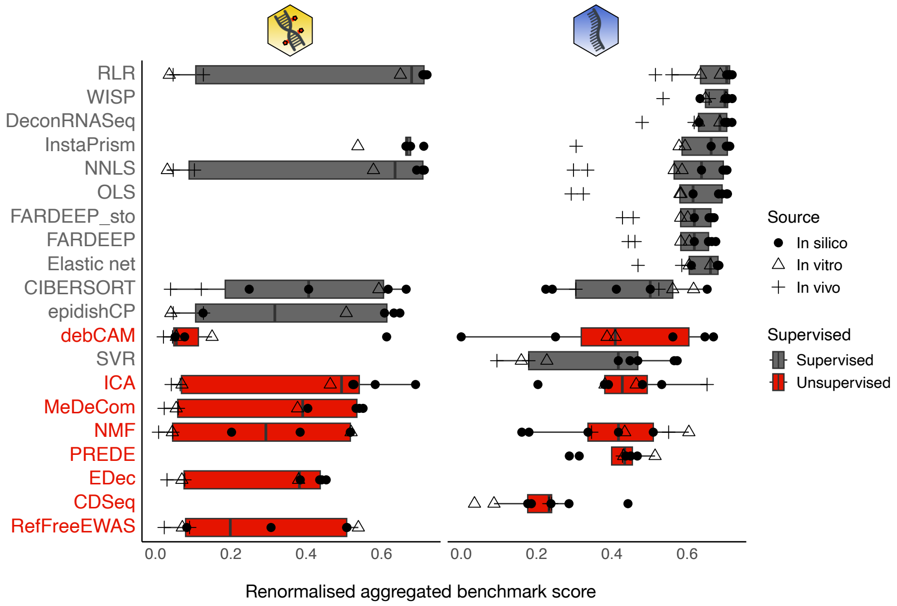


**Supplementary Figure 12** Renormalized aggregated benchmark scores. Candidates are ordered from top to bottom by their overall benchmark score. The yellow DNA icon represents DNAm methods, the blue single-strand one RNA methods.

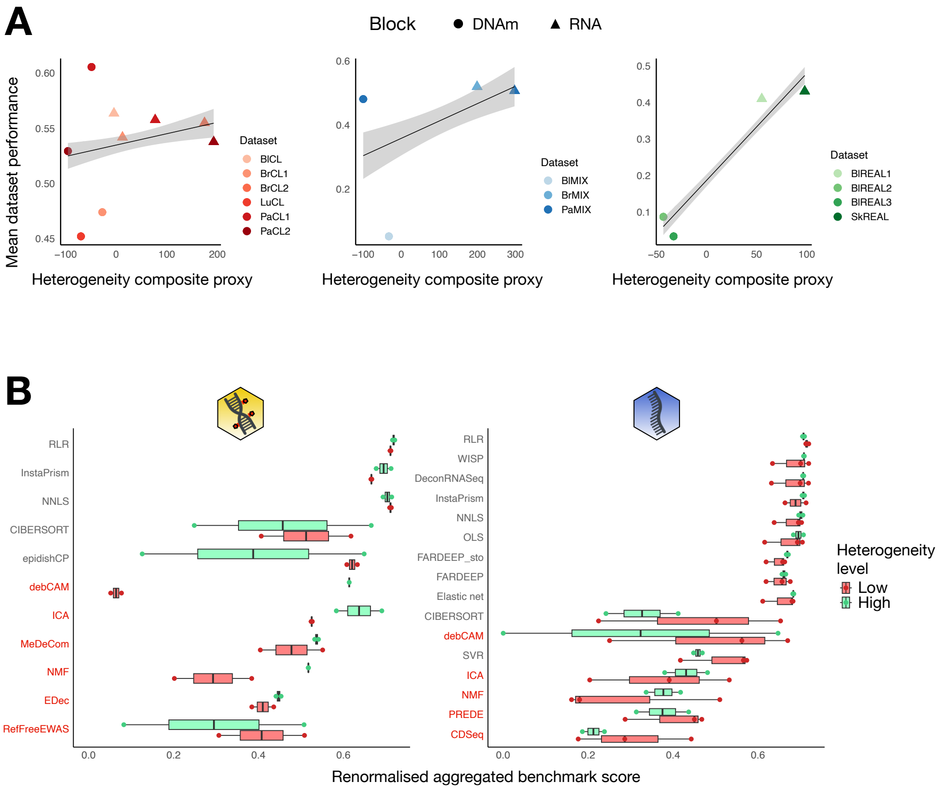


**Supplementary Figure 13** Impact of heterogeneity levels on the general deconvolution performance. **(A)**Global deconvolution performance is weakly impacted by dataset heterogeneity. Each facet represents a data source: *in silico*, *in vitro* and *in vivo*. **(B)** Effect of the heterogeneity level on each method's performance as a function of the omic type: the yellow icon stands for DNAm methods, the blue one for RNA methods. Unsupervised methods are in red.

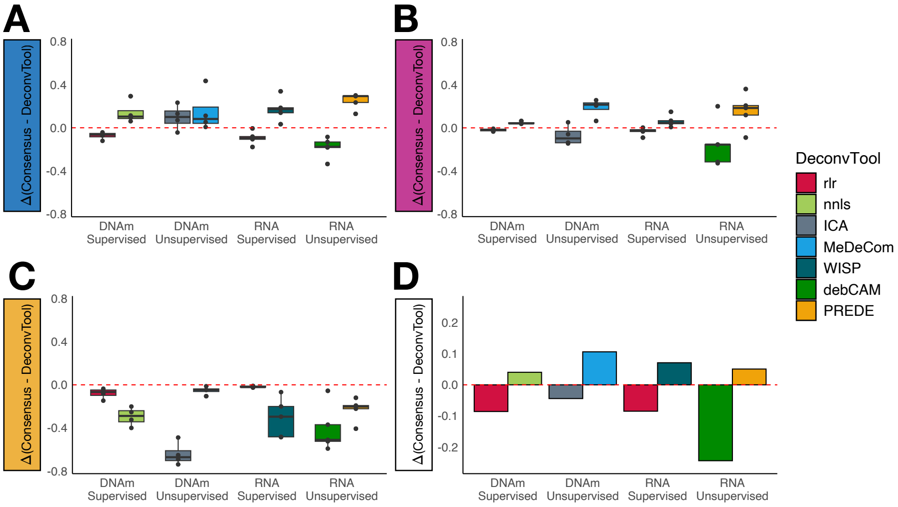


**Supplementary Figure 14** Consensus approaches. They do not improve the performance of the deconvolution, in any combination of class and omic. The boxplots display on the y-axis the differences in normalised scores for each dataset between the consensus strategy and the two best methods in each setting. The raw performance is shown in panel **(A)**, the stability in panel **(B)** and the scalability in panel **(C)**. **(D)** Difference in overall score between the consensus strategy and the two best methods of each setting.

| **Method** | **Approach** | **DOI** | **Reviewed in** | **Supervised** | **Language** | **Included** | **Details** |
|---|---|---|---|---|---|---|---|
| **ARIC** | $\nu$-SVR | 10.1093/bib/bbab362 | 1, 2 | Yes | Python | No | |
| **BayesCCE** | Bayesian | 10.1186/s13059-018-1513-2 | 2, 3 | Semi | MATLAB | No | |
| **CelFEER** | EM | 10.1093/nargab/lqad048 | 2 | Semi | Python | No | |
| **CelFiE** | EM | 10.1038/s41467-021-22901-x | 2 | Semi | Python | No | |
| **cfSort** | DNN | 10.1073/pnas.2305236120 | 2 | Yes | Python | No | |
| **CIBERSORT** | $\nu$-SVR | 10.1038/nmeth.3337 | 2, 4, 5, 7 | Yes | R, Web-based | Yes | |
| **CIBERSORTx** | $\nu$-SVR | 10.1038/s41587-019-0114-2 | 2 | Yes | Web-based | No | |
| **DXM** | HMM | 10.1093/nar/gkab516 | 2, 6 | No | Python | No | |
| **EDec** | NMF | 10.1016/j.celrep.2016.10.057 | 2, 3 | No | R | Yes | |
| **Emeth** | EM | 10.1038/s41598-021-84864-9 | 2 | Semi | R | No | |
| **epidishCP** | CLS | 10.1186/1471-2105-13-86 | 1, 2, 3, 4, 5, 6, 7 | Yes | R, Python | Yes | |
| **EPISCORE** | RPC-LS | 10.1186/s13059-020-02126-9 | 2, 3 | Yes | R | No | Requires scRNAseq data to impute a DNAm reference matrix |
| **HiBED** | HM | 10.3389/fnins.2023.1198243 | 2 | Yes | R | No | Specific to cerebral data |
| **HiTIMED** | HM | 10.1186/s12967-022-03736-6 | 2 | Yes | R | No | Similar to epidishCP |
| **ICA** | ICA | 10.3390/ijms20184414 | 3 | No | R, Python | Yes | |
| **MeDeCom** | NMF | 0.1186/s13059-017-1182-6 | 2, 3, 6 | No | R | Yes | |
| **MethAtlas** | NNLS | 10.1038/s41467-018-07466-6 | 2 | Yes | Python | No | |
| **MethylPurify** | EM | 10.1186/s13059-014-0419-x | 2, 6 | No | Python | No | |
| **MethylResolver** | LS | 10.1038/s42003-020-01146-2 | 1, 2, 3 | Yes | R | No | Designed only for immune cells |
| **NMF/RefFreeCellMix** | NMF | 10.1186/s12859-016-1140-4 | 2, 3, 4, 5, 7 | No | R | Yes | |
| **PRISM** | EM | 10.1093/bioinformatics/btz327 | 2, 6 | No | Python | No | |
| **PRMeth** | NMF | 10.1186/s12859-022-04893-7 | 2 | Semi | R | No | |
| **RefFreeEWAS** | NMF | 10.1093/bioinformatics/btu029 | 1, 2, 4, 5, 7 | No | R | Yes | |
| **RLR** | RPC-LS | 10.1186/s12859-017-1511-5 | 1, 2, 3, 4, 5, 7 | Yes | R | Yes | |
| **Tsisal** | Geometric | 10.1093/bioinformatics/btaa930 | 2 | Both | R | No | |
| **UXM** | NNLS | 10.1038/s41586-022-05580-6 | 2 | Yes | Python | No | |

**Supplementary Table 1** Literature review of methylation deconvolution methods. Methods in green are those who were included in this article. References for the reviews listed in the table are in the Supplementary Methods. SVR: Support Vector Regression. EM: Expectation–Maximization. DNN: Deep Neural Network. HMM: Hidden Markov model. NMF: Non-negative Matrix Factorization. LS: Least Squares. CLS: Constrained LS. RPC: Robust Partial Correlation. HM: Hierarchical Model. ICA: Independent Component Analysis. NNLS: Non Negative LS.

| Method | Approach | DOI | Reviewed in | Supervised | Language | Included | Details |
|---|---|---|---|---|---|---|---|
| **Abbas** | CLS | 10.1371/journal.pone.0006098 | 1, 2 | Yes | R | No | Similar to DeconRNASeq |
| **AutogeneS** | $\nu$-SVR | 10.1016/j.cels.2021.05.006 | 3, 4 | Yes | Python | No | |
| **BayesPrism/InstaPrism** | Bayesian | 10.1038/s43018-022-00356-3 | 3, 4, 5 | Yes | R | Yes | |
| **BisqueRNA** | CLS | 10.1038/s41467-020-15816-6 | 3, 4, 5, 6 | Yes | R | No | Relies on single-cell reference |
| **Bseq-SC** | $\nu$-SVR | 10.1016/j.cels.2016.08.011 | 3, 4 | Yes | R | No | Relies on single-cell reference |
| **CDSeq** | LDA | 10.1371/journal.pcbi.1007510 | 4 | No | R | Yes | |
| **CIBERSORT/SVR** | $\nu$-SVR | 10.1038/nmeth.3337 | 2, 3, 4, 6, 7, 8, 9, 10 | Yes | R | Yes | |
| **CIBERSORTx** | $\nu$-SVR | 10.1038/s41587-019-0114-2 | 4, 5, 7, 8 | Yes | Web-based | No | |
| **CPM** | SVR | 10.1038/s41592-019-0355-5 | 3, 4, 5 | Yes | R | No | Relies on single-cell reference |
| **debCAM** | Convex | 10.1038/srep18909 | 3, 4, 7 | No | R | Yes | |
| **deconf** | NMF | 10.1186/1471-2105-11-27 | 2, 3, 4 | No | R | No | Developed for microarrays |
| **DeconRNASeq** | CLS | 10.1093/bioinformatics/btt090 | 2, 3, 4, 6, 7, 9 | Yes | R | Yes | |
| **DSection** | Bayesian | 10.1093/bioinformatics/btq406 | 1, 11 | No | MATLAB | No | |
| **dtangle** | LS | 10.1093/bioinformatics/bty926 | 3, 4, 6, 9 | Yes | R | No | Relies on markers |
| **DWLS** | CLS | 10.1038/s41467-019-10802-z | 3, 4, 5, 6 | Yes | R | No | Relies on single-cell reference |
| **Elastic net** | LS | 10.18637/jss.v033.i01 | 4, 6 | Yes | R | Yes | |
| **EPIC** | CLS | 10.7554/eLife.26476 | 2, 3, 4, 5, 6, 7, 8, 10 | Yes | R | | |
| **FARDEEP** | CLS | 10.1371/journal.pcbi.1006976 | 3, 4, 6 | Yes | R | Yes | |
| **ICA** | ICA | 10.3390/ijms20184414 | 3 | No | R | Yes | |
| **ImmuCellAI** | CLS | 10.1002/advs.201902880 | 3 | Yes | R | | |
| **Lasso** | LS | 10.18637/jss.v033.i01 | 4, 6 | Yes | R | | |
| **LinDeconSeq** | CLS | 10.1186/s12864-020-06888-1 | 3 | Yes | R | No | Relies on single-cell reference |
| **LinSeed** | Linear | 10.1038/s41467-019-09990-5 | 3, 7, 9 | No | R | No | Relies on markers |
| **MCP-Counter** | Linear | 10.1186/s13059-016-1070-5 | 2, 3, 8, 10 | Yes | R | No | Relies on markers |
| **MMAD** | MLE | 10.1093/bioinformatics/btt566 | 2, 7 | No | MATLAB | No | |
| **MOMF** | NMF | 10.5281/zenodo.3373980 | 3, 4 | Yes | R | No | Relies on single-cell reference |
| **MuSiC** | CLS | 10.1038/s41467-018-08023-x | 3, 4, 5, 6, 7, 9 | Yes | R | No | Relies on single-cell reference |
| **NITUMID** | NMF | 10.1093/bioinformatics/btz748 | 3 | Yes | R | | |
| **NNLS** | CLS | 10.1137/1.9781611971217 | 4, 6 | Yes | R | Yes | |
| **OLS** | LS | 10.1007/978-3-642-50096-1_48 | 4, 6 | Yes | R | Yes | |
| **PERT** | MLE | 10.1371/journal.pcbi.1002838 | 1, 2 | Yes | Octave | No | |
| **PREDE** | NMF | 10.1371/journal.pcbi.1008452 | 3 | Yes | R | Yes | |
| **proportionsInAdmixture** | LS | 10.1371/journal.pone.0224693 | 4 | Yes | | | |
| **quanTIseq** | CLS | 10.1186/s13073-019-0638-6 | 2, 3, 8, 10 | Yes | R/Docker | | |
| **Ridge/AdRoit** | CLS | 10.18637/jss.v033.i01 | 3, 4, 6 | Yes | R | No | Relies on single-cell reference |
| **RLR/EpiDISH** | RPC-LS | 10.1038/s41592-018-0213-x | 4, 6 | Yes | R | Yes | |
| **SCDC** | Ensemble | 10.1093/bib/bbz166 | 3, 4, 6 | Yes | R | No | Relies on single-cell reference |
| **ssFrobenius** | NMF | 10.1186/1471-2105-11-27 | 2, 4, 6 | Semi | R | No | |
| **ssKL** | NMF | 10.1186/1471-2105-11-27 | 2, 4, 6 | Semi | R | No | |
| **ssNMF** | NMF | 10.1016/j.meegid.2011.08.014 | 1 | Semi | R | No | |
| **TIMER** | CLS | 10.1186/s13059-016-1028-7 | 2, 4, 7, 10 | Yes | Web-based | No | |
| **WISP** | CLS | 10.1038/s41467-019-09307-6 | | Yes | R | Yes | |
| **xCell** | GSEA | 10.1186/s13059-017-1349-1 | 2, 8, 9, 10 | Yes | R | No | Computes enrichment |

**Supplementary Table 2** Literature review of transcriptome deconvolution methods. Methods in green are those who were included in this article. References for the reviews listed in the table are in the Supplementary Methods. LS: Least Squares. CLS: Constrained LS. SVR: Support Vector Regression. LDA: Latent Dirichlet Allocation. NMF: Non-negative Matrix Factorization. MLE: Maximum Likelihood Estimation. ICA: Independent Component Analysis. PCA: Principal Component Analysis. RPC: Robust Partial Correlation. GSEA: Gene Set Enrichment Analysis.

| Dataset | Fibroblast | T cell | B cell | NK cell | Monocyte | Neutrophil | Immune cell | Breast | Lung | Pancreas |
|---|---|---|---|---|---|---|---|---|---|---|
| **BrCL1** | 45% | 10% | | | | | | 45% (15% healthy, 30% cancer) | | |
| **PaCL1** | 45% | | | | | | 10% | | | 45(% (15% endothelial, 30% cancer) |
| **PaCL2** | 46% | 7% (4% CD4$^+$, 3% CD8$^+$) | 1% | | | 1% | 1% (Macrophage) | | | 44% (15% endothelial, 29% cancer) |
| **BlCL** | | 60% | 13% | 7% | 15% | 4% | 1% (mDC) | | | |
| **BrCL2** | 45% | 5% (CD4$^+$) | | | 5% | | | 45% (15% BT474, 15% MCF7, 15% T47D) | | |
| **LuCL** | | 10% (5% CD4$^+$, 5% CD8$^+$) | 4% | 4% | 4% | 4% | 4% (WBC) | | 70% (10% NHBEC, 60% A549) | |

**Supplementary Table 3** Proportions $\alpha_i$ used for the different cell types in the simulated datasets. Rare cell types ($\alpha_i < 5\%$) are in green.
