## Supplementary Methods for "A robust workflow to benchmark deconvolution of multi-omic data"

### A ROBUST WORKFLOW TO BENCHMARK DECONVOLUTION OF MULTI-OMIC DATA
### SUPPLEMENTARY METHODS

#### Data

##### Simulations: model technical noise

We added a noise after convoluting the reference matrix $T$ with the proportions matrix $A$ to mimic the technical noise.

**Technical noise for DNAm data** For methylation data, we added a Gaussian noise with parameters $\mu = 0$, $\sigma = 3$. More precisely, we did the convolution $T \times A$, and converted the resulting matrix to M-values, using the formula $M = log2(\beta)/(1-\beta)$. Then we generated random numbers based on a Gaussian distribution with the *rnorm* R function, that we added to the M-values matrix. Finally, we revert to $\beta$-values using the formula $\beta = 2^M/(2^M+1)$.

**Technical noise for RNA data** For transcriptomic data, we used a Negative Binomial noise. We designed the following procedure to generate a Negative Binomial Noise to the convolution $d = T \times A$:

$p = 0.1$
$\sigma = 1$
$\Delta$ is a matrix of Gaussianly distributed numbers with the parameters $\mu = 0$ and $\sigma$, of the same size as $d$
$c$ is the vector of the sums on the columns of $d$
$\mu_0 = \frac{d}{c} \times \text{mean}(c)$
$\sigma_0 = (1.8p + \frac{1}{\sqrt{\mu_0}}) \times \exp(\frac{\Delta}{2})$
shape $= \frac{1}{\sigma_0^2}$
scale $= \frac{\mu_0}{\text{shape}+\epsilon}$
$\mu$ is a matrix of numbers drawn from a Gamma distribution with parameters shape and scale of the same size as $d$
The resulting expression matrix $D$ is a matrix of numbers drawn from a Poisson distribution with parameters $\mu$

##### Creating an extra cell type

To create an extra cell type in the reference matrix, we first started by selecting all immune cell types $T_{\text{immune}}$ in $T$. If we have more than one cell type, we computed the average profile by calculating the arithmetic mean for each row of $T_{\text{immune}}$. Finally we added noise, following the procedure described above. For methylation data, the parameters we used for the Gaussian distribution were $\mu = 0$ and $\sigma = 1$. For transcriptomic data, the parameters $p$ and $\sigma$ were kept the same.

##### Quantifying the heterogeneity of a dataset

We used two proxies to quantify a dataset's heterogeneity. First, we measured the mean of the Pearson correlations between all pairs of cell types in $T$, called "Mean $R^2$". Then, we computed the logarithm of the phenotypic volume $V$ as defined in.[1] Briefly, it represents the volume occupied by the different cell types in the space of molecular features:

$$\log(V) = \log(\frac{|\Sigma|_+}{K-1}) = \log(\frac{\prod_{e=1}^{E} \lambda_e}{K-1}); \forall \lambda_e > 0$$

where $\Sigma$ is the covariance matrix of $T$ restricted to the $K-1$ most variable features, $K$ is the number of cell types and $\lambda_e$ are the non-zero eigenvalues of $\Sigma$.

We used in our analyses the composite parameter HG $= \log(V) \times R^{-2}$.

#### Deconvolution pipeline: methods

The RDS file containing genes' length for TPM normalization is available upon request to the corresponding authors. We specify below the functions and the non-default parameters that were used for each deconvolution method. As a reminder, NN stands for Non Negative and STO for Sum To One.

##### RNA methods

CIBERSORT was run with the *EpiDISH::epidish* function, setting "method" to "CBS".

DeconRNASeq was run with the *DeconRNASeq::DeconRNASeq* function with "use.scale = FALSE" and a further STO constraint.

Elastic net was done via the *glmnet::glmnet* function with further explicit NN and STO constraints.

FARDEEP and FARDEEP_sto were run with the *FARDEEP::fardeep* function, and proportions were retrieved from the "abs.beta" slot for FARDEEP, and "relative.beta" slot for FARDEEP_sto.

InstaPrism was run by adapting the code available at https://github.com/humengying0907/InstaPrism/tree/master, and setting the number of cores to 32.

NNLS, OLS and SVR were run with the *granulator* package on TPM-normalized data with 32 cores, with further explicit NN and STO constraints.

RLR was done with the *EpiDISH::epidish* function, setting "method" to "RPC".

WISP was run using the code available on https://github.com/cit-bioinfo/WISP, adding an additional STO constraint.

CDSeq, PREDE and debCAM were run with the same parameters as in https://github.com/bcm-uga/gedepir/blob/main/R/run_deconvolution.R, with a NN constraint for PREDE.

ICA was run with the *fastICA::fastICA* function with "maxit = 1000" and "tol = $1 \times 10^{-9}$", with a further STO constraint.

NMF with performed with the *NMF::nmf* function on the matrix restricted to the features that have at least 1 count across all the samples, with a seed of 1 and method = "snmf/r", and a STO constraint.

##### DNAm methods

CIBERSORT, NNLS and RLR were run as described for RNA methods.

epidishCP was run with the *EpiDISH::epidish* function, setting "method" to "CP".

debCAM, ICA and NMF were run as described above.

EDec was performed with the *EDec::run_edec_stage_1* function using all CpGs as informative loci, and with an additional NN constraint.

InstaPrism was run as described above, with a pre-transformation of the data by multiplying the $\beta$-values by 1,000 and rounding the result.

RefFreeEWAS was run with the *RefFreeEWAS::RefFreeCellMix* function and additional NN and STO constraints.

**References of the reviews cited in Supplementary Tables 1 and 2**

| Review number | DOI |
|---|---|
| 1 | 10.1093/bib/bbac449 |
| 2 | 10.1093/bib/bbae234 |
| 3 | 10.1038/s43588-021-00038-7 |
| 4 | 10.2217/epi-2016-0153 |
| 5 | 10.1093/hmg/ddx275 |
| 6 | 10.1093/bib/bbac248 |
| 7 | 10.1016/j.csbj.2021.12.001 |

Reviews listed in Supplementary Table 1

| Review number | DOI |
|---|---|
| 1 | 10.1016/j.coi.2013.09.015 |
| 2 | 10.1007/s00262-018-2150-z |
| 3 | 10.1093/nar/gkae267 |
| 4 | 10.1093/bioadv/vbae048 |
| 5 | 10.1038/s41467-023-41385-5 |
| 6 | 10.1038/s41467-020-19015-1 |
| 7 | 10.1186/s13059-021-02290-6 |
| 8 | 10.1038/s41467-024-50618-0 |
| 9 | 10.1038/s41467-022-28655-4 |
| 10 | 10.1093/bioinformatics/btz363 |
| 11 | 10.1093/bib/bbu002 |

Reviews listed in Supplementary Table 2

#### Ranking pipeline

##### Metrics preparation

We display in Table 1 an example of the primary metrics we can compute directly from the estimation of the proportions by a given method for each simulation. Of note, the time $t$ is the real execution time divided by the number of samples. In case we have replicates, *i.e.* for *in silico* data, we can add secondary stability metrics, and we average across replicates for the primary metrics, as in Table 2. The time is also log-transformed since the execution times of all methods spanned several orders of magnitude:

$\mathrm{Time}_{\mathrm{log}} = \log_{10}(1 + \mathrm{t})$

| Replicate | RMSE | MAE | Pearson_Matrix | Pearson_CellType | Pearson_Sample | Time (s) |
|---|---|---|---|---|---|---|
| Sim1 | 0.21 | 0.33 | 0.87 | Median = 0.70<br>Standard dev = 0.02 | Median = 0.87<br>Standard dev = 0.03 | 12 |
| Sim2 | 0.23 | 0.27 | 0.92 | Median = 0.79<br>Standard dev = 0.01 | Median = 0.85<br>Standard dev = 0.01 | 11 |
| Sim3 | 0.18 | 0.40 | 0.86 | Median = 0.83<br>Standard dev = 0.04 | Median = 0.89<br>Standard dev = 0.02 | 13 |
| Sim4 | 0.20 | 0.41 | 0.80 | Median = 0.81<br>Standard dev = 0.03 | Median = 0.90<br>Standard dev = 0.02 | 12 |

Table 1: Fake example of primary metrics.

Then we normalize the different metrics. We scale all metrics between 0 and 1, and we transform the scores such that 1 is always the best result, by inverting all metrics except the Pearson ones. For the normalization step, we first center all scores from all methods for a given metric and a given dataset by subtracting the arithmetic mean. Then we scale by dividing by the standard deviation and applying the distribution function (with the *pnorm* R function) of the normal distribution.

| RMSE | MAE | Pearson_Matrix | Pearson_CellType | Pearson_Sample | Time (log) |
|---|---|---|---|---|---|
| Median = 0.21<br>Sd = 0.02 | Median = 0.37<br>Sd = 0.07 | Median = 0.87<br>Sd = 0.05 | Median of medians = 0.80<br>Median of standard devs = 0.03 | Median of medians = 0.88<br>Median of standard devs = 0.02 | Median = 1.11<br>Sd = 0.03 |

Table 2: Fake example of averaged primary metrics along with secondary metrics.

The last step is to merge Pearson metrics into a single Pearson meta-score, separately for the metrics related to raw performance or to stability. We select first the averaged-normalized-transformed Pearson_Matrix, Pearson_CellType and Pearson_Sample metrics and calculate the arithmetic mean of those three scores for each method and dataset. We do the same for the standard deviation of the normalized-transformed Pearson_Matrix, Pearson_CellType and Pearson_Sample metrics. We end up with 8 metrics in case of replicates (4 otherwise), as shown in Table 3.

| RMSE | MAE | Pearson | Time (log) |
|---|---|---|---|
| Norm median = 0.79<br>Norm Sd = 0.98 | Norm median = 0.63<br>Norm sd = 0.93 | Norm mean = 0.85<br>Norm sd = 0.97 | Norm median = 0.82<br>Sd = 0.97 |

Table 3: Fake example of averaged-normalized-transformed scores after merging Pearson metrics.

##### Metrics renormalization to measure a dataset's easiness-of-deconvolution

Since we wanted to quantify how easy it was to deconvolve a specific dataset, we do not want to normalize scores as previously described. Indeed, this procedure is meant to avoid having a dataset with a large impact on the overall score by smoothing out differences between ease- and hard-to-deconvolve datasets. Here, we re-normalized each metric across all methods and all datasets simultaneously, instead of normalizing on a per dataset basis. Then we transformed the scores and merged all Pearson metrics. Finally, we computed the mean of the re-normalized aggregated scores for all methods for a dataset $X$ to get a metric measuring dataset's easiness of deconvolution.

##### Metrics aggregation

The global aggregation procedure is broken down in three steps. The first step (i) consists in merging all metrics inside a category (raw performance, stability or scalability) and for a given dataset and method with a geometric mean as it is less influenced by outlier metrics than the arithmetic mean. We used the geometric mean whenever we wanted to mitigate the presence of possible outliers. The second step (ii) consists in merging, for a given dataset and method, the three categories with a weighted geometric mean. We used a weight of 1 for the raw performance, which we deemed to be the most important parameter to consider, and weights of 0.5 for stability and scalability. The last step (iii) is the final merging of the scores for one method for the different datasets with an arithmetic mean. The whole procedure is summarized in scheme 1.

##### Process $S_{rank}$

We have a ranking process that relies on computing ranks from the normalized-transformed scores before doing the global aggregation. For a fixed dataset and metric, we calculate the ranks of all methods with the R function *rank*, with the parameter "ties.method" set to "average" and the parameter "na.last" set to "keep". We then scale the ranks between 0 and 1 by dividing by the maximum rank, before going through the global aggregation described above.

##### Process $S_{topsis}$

For the topsis process, we use the normalized-transformed scores to compute TOPSIS scores that quantify how close a method is to the positive ideal solution (PIS) and how far it is from the negative ideal solution (NIS). In practice, a TOPSIS score for a given method and metric in a fixed dataset is calculated as follows: the PIS is defined as the maximum score observed across all methods for each metric, and the NIS as the minimum score. Then we calculate for each metric the Euclidean distance from a method to the PIS and NIS. Finally the TOPSIS score is the ratio $\frac{d_{\text{NIS}}}{d_{\text{NIS}}+d_{\text{PIS}}}$. Hence the TOPSIS score is restricted to the $[0, 1]$ interval and the higher it is, the closer the method is to the PIS. From the matrix of TOPSIS scores, we do the global aggregation to perform the process $S_{topsis}$.

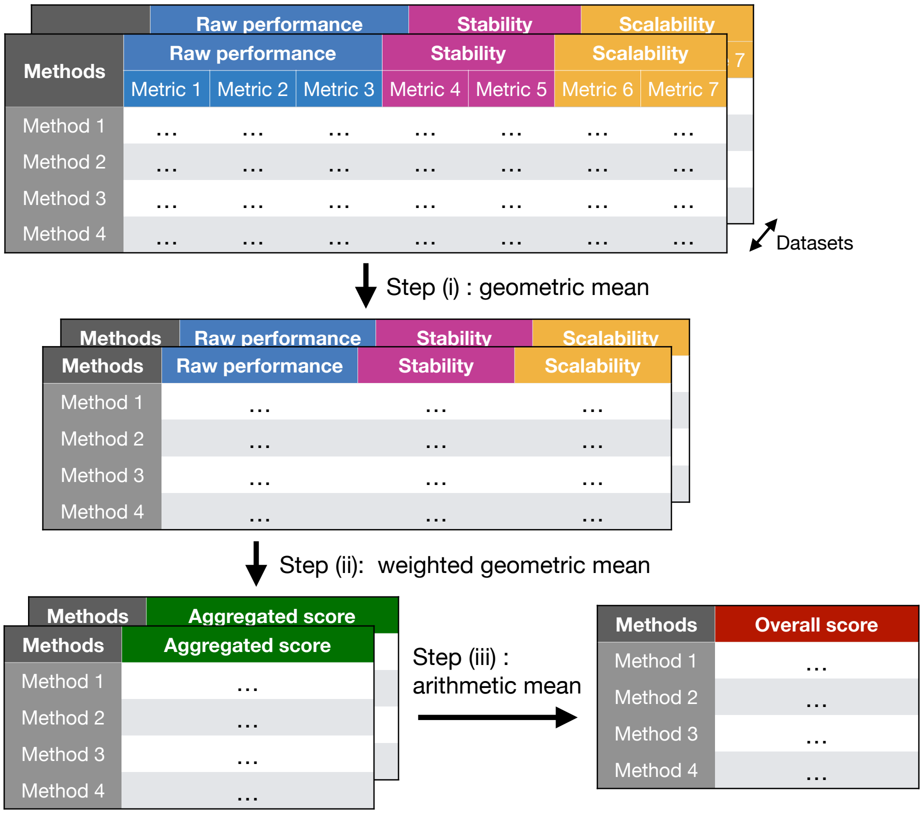


Scheme 1: Scheme of the three-steps global aggregation procedure.

##### Criteria for the evaluation of ranking processes

We were interested in evaluating the soundness of the ranking processes we designed. We quantified several empirical criteria as in:[2] the average rank, the condorcet rate and the generalization criterion. Let us briefly describe these, each one starting with the matrix of normalized-transformed scores $\mathbf{M}$ with the methods on the rows and the metrics on the columns. We calculated each criterion for each setting, which is a combination of the omic analysed and the class of the methods, supervised or unsupervised.

**Average rank** The average rank looks at the mean rank of the winner, across all metrics, elected by the ranking process $S$ being evaluated. If $x$ is the winner method, we first compute the matrix of ranks across all metrics $\mathbf{R}^{\mathbf{M}}$, then the mean rank of the winner $mean(\mathbf{R}^{\mathbf{M}}{}_{x,})$. Finally we normalize the result to obtain a value between 0 and 1, 1 being the best score:

$1 - \frac{mean(\mathbf{R}^{\mathbf{M}}{}_{x,})-1}{J-1}$

with $J$ the number of metrics.

**Condorcet rate** Let us first define the Condorcet winner. The Condorcet winner is the method that would win across more than 50% of the metrics against all the other methods in a pairwise comparison. Hence the Condorcet rate is the rate at which the winner elected by our ranking process corresponds to the Condorcet winner in all pairwise comparisons.

**Generalization criterion** This criterion quantifies how generalizable the ranking process is, meaning how much it depends on a subset of metrics or not. The higher the criterion, the less it depends on specific metrics. It is intended to measure how much adding new metrics will perturb the ranking. Shortly, it is calculated for each ranking process as follows:

Gen $= \frac{1}{|\mathcal{J}^{test}|} \sum_{j \in \mathcal{J}^{test}} \sigma(\mathbf{S}^{train}, \mathbf{S}^{test})$

where $\sigma$ is the Spearman correlation, $\mathcal{J}^{train}$ and $\mathcal{J}^{test}$ are disjoint sets of metrics, and $\mathbf{S}^{train}$ (resp. $\mathbf{S}^{test}$) are the ranks obtained from applying the ranking process $S$ on the matrix $\mathbf{M}$ restricted to the metrics in $\mathcal{J}^{train}$ (resp. $\mathcal{J}^{test}$).

In our case, we excluded 10% of the metrics in the train set. Since we don't have many metrics, it led to having all metrics except one in the train set, and one metric in the test set, hence $|\mathcal{J}^{test}|= 1$. We excluded successively the first, second, ..., k-th, ..., $J$-th metric and computed the corresponding value $\mathsf{Gen}_k$:

$\mathsf{Gen}_k = \sigma(\mathbf{S}^{-k}, \mathbf{S}^k)$

where $\mathbf{S}^{-k}$ (resp. $\mathbf{S}^k$) are the ranks obtained using all the metrics minus the k-th one (resp. using the k-th metric).

Finally, we get:

$\mathsf{Gen} = \frac{1}{J}\sum_{k=1}^{J} \mathsf{Gen}_k$

### References


[1] Elham Azizi, Ambrose J. Carr, George Plitas, Andrew E. Cornish, Catherine Konopacki, Sandhya Prabhakaran, Juozas Nainys, Kenmin Wu, Vaidotas Kiseliovas, Manu Setty, Kristy Choi, Rachel M. Fromme, Phuong Dao, Peter T. McKenney, Ruby C. Wasti, Krishna Kadaveru, Linas Mazutis, Alexander Y. Rudensky, and Dana Pe'er. Single-Cell Map of Diverse Immune Phenotypes in the Breast Tumor Microenvironment. *Cell*, 174(5):1293–1308.e36, August 2018.

[2] Adrien Pavao, Michael Vaccaro, and Isabelle Guyon. Judging competitions and benchmarks: a candidate election approach, 2021. Paper presented at ESANN 2021.
